# UHRF1 restricts HCoV-229E infection through epigenetic silencing of the viral receptor APN

**DOI:** 10.1101/2025.08.19.671085

**Authors:** Pengcheng Wang, Ziqiao Wang, Fei Feng, Lisha Yin, Yuyuan Zhang, Zhichao Gao, Jiannan Chen, Ping Zhang, Shuiqiao Yuan, Qiang Ding, Yue Hong, Yuanlin Song, Chun Li, Jincun Zhao, Rong Zhang

## Abstract

The emergence of SARS-CoV-2 has posed significant threats to global health, particularly for the older population. Similarly, common human coronaviruses, such as HCoV-229E, which typically cause mild cold-like symptoms, have also been linked to severe diseases, underscoring the need to understand virus-host interactions and identify host factors contributing to viral pathogenesis and disease progression. In this study, we performed a genome-wide CRISPR knockout screen using HCoV-229E and identified Ubiquitin-like with PHD and RING finger domain 1 (UHRF1) as a potent restriction factor. Mechanistically, UHRF1 suppressed HCoV-229E infection by downregulating the expression of its cell entry receptor, aminopeptidase N (APN), through promoter hypermethylation. Focused CRISPR activation screens of UHRF1-downregulated genes confirmed the critical role of APN in HCoV-229E infection and identified additional genes (e.g., SIGLEC1, PLAC8, and heparan sulfate biosynthesis genes) contributing to the restrictive functions of UHRF1. Transcriptomic and single-cell RNA sequencing analysis revealed that UHRF1 expression decreases with age, negatively correlating with increased APN expression. This age-related decline in UHRF1 was further validated in primary alveolar macrophages isolated from elderly individuals, which exhibited heightened susceptibility to HCoV-229E infection compared to those from younger individuals. Our findings highlight UHRF1 as a key age-related host defense factor against coronavirus infection and provide novel insights into the epigenetic regulation of viral entry receptors.

## INTRODUCTION

The threat posed by the COVID-19 pandemic has gradually diminished with the development of vaccines and antiviral drugs. In contrast to SARS-CoV-2 and two other coronaviruses (MERS-CoV and SARS-CoV), which cause severe diseases with high morbidity and mortality, common human coronaviruses (e.g., HCoV-OC43, HCoV-229E, HCoV-NL63, and HCoV-HKU1) typically induce mild, cold-like symptoms^1,2^. Among these human coronaviruses with low pathogenicity, HCoV-229E, a member of the genus *Alphacoronavirus*, is associated with mild, self-limited symptoms such as runny nose, sore throat, headache, cough, and fever^3^. However, HCoV-229E has also been linked with lower respiratory tract infections, including life-threatening pneumonia, bronchiolitis, and acute respiratory distress syndrome, particularly in immunodeficient and elderly patients^4–6^.

Notably, HCoV-229E can efficiently invade the primary (alveolar) macrophages and monocytes with productive infection, unlike other coronaviruses^7–10^. Infection of myeloid cells by HCoV-229E may facilitate viral dissemination to other tissues, potentially inducing systemic pathologies. Alveolar macrophages are essential lung resident cells, serving as the first line of defense in maintaining homeostasis, combating pathogens, and regulating lung inflammation^7,11^. Human aminopeptidase N (APN), also known as CD13 or alanyl aminopeptidase, functions as the cell entry receptor for HCoV-229E^12^. The expression of APN in (alveolar) macrophages and monocytes likely explains their susceptibility to HCoV-229E infection.

Viruses rely heavily on host factors for their replication and pathogenesis. Identifying these factors is of significance for advancing our understanding of host-pathogen interactions. While host dependency factors that promote viral infection are important to explore, host restriction factors that modulate viral tropism and pathogenesis should not be overlooked. To identify host restriction factors associated with susceptibility to HCoV-229E and disease severity, we performed a genome-wide CRISPR knockout screen and found the Ubiquitin-like with PHD and RING finger domain 1 (UHRF1), also known as inverted CCAAT box-binding protein of 90 kDa (ICBP90), as the top-ranked candidate.

UHRF1 plays an important role in maintaining DNA methylation by recruiting DNA methyltransferase I (DNMT1) to hemi-methylated DNA^13^ and is involved in various biological processes, including spermatogenesis, tumor progression, and cellular metabolism^14–17^. UHRF1 also influences infection by multiple viruses (e.g., human immunodeficiency virus 1 [HIV-1], Epstein-Barr virus [EBV], alphaherpesviruses, influenza virus, and vesicular stomatitis virus [VSV])^18–22^. However, its role in coronavirus infection has not been previously documented. Here, we demonstrate for the first time that UHRF1 restricts HCoV-229E infection primarily by suppressing the expression of its receptor APN, through promoter methylation. Furthermore, we reveal a negative correlation between UHRF1 expression and age, resulting in increased susceptibility of primary alveolar macrophages from elderly individuals to HCoV-229E. These findings enhance our understanding of virus-pathogen interactions and provide insights into the development of antiviral strategies against coronaviruses such as HCoV-229E.

## RESULTS

### Genome-wide CRISPR knockout screen identifies UHRF1 as a host restriction factor for HCoV-229E infection

To identify host intrinsic factors capable of restricting coronavirus infection, particularly HCoV-229E, which causes common cold-like symptoms but can also lead to severe disease in elderly patients, we performed a genome-wide CRISPR knockout screen using an A549 cell library. We used the HCoV-229E expressing the mGreenLantern reporter in place of the ns4a gene (HCoV-229E-mGreen) as a model virus. Given the inefficiency of HCoV-229E infection in A549 cells, we hypothesized that host restriction factors could be identified by sorting susceptible cells following gene knockout. At 24 hours post-infection, reporter-positive infected cells were collected by flow cytometry for DNA extraction and next-generation sequencing. After data analysis **(Supplementary Table 1)**, we identified top candidates based on MAGeCK scores, P-values, and false discovery rate (FDR) **(Fig. 1A and Supplementary Fig. 1)**, with *UHRF1* ranking as the top candidate.

**Figure 1.**
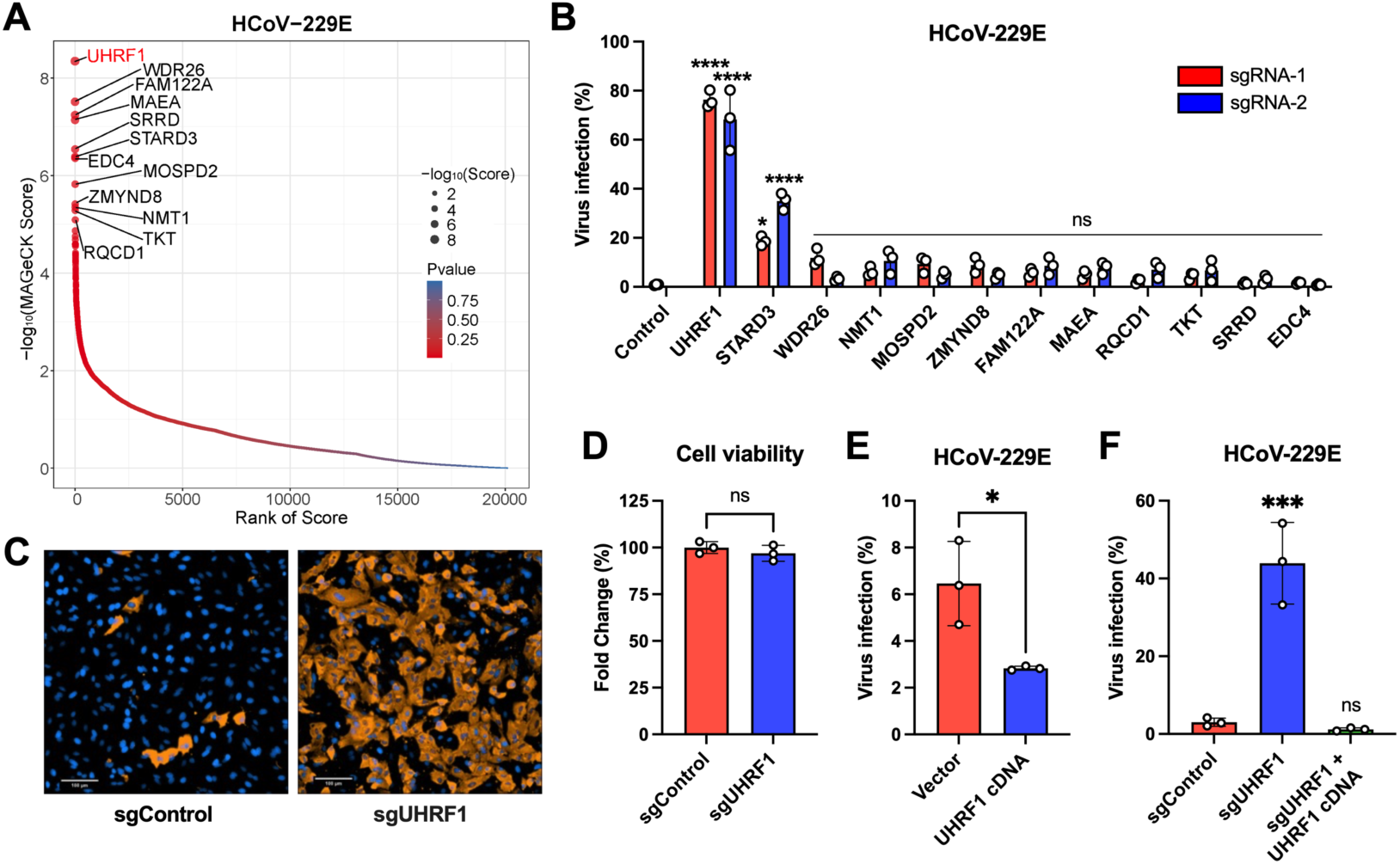
Genome-wide CRISPR knockout screen identifies UHRF1 as a host restriction factor for HCoV-229E infection. **A.** Identification of host restriction genes from the CRISPR knockout screen. The A549 cell library containing genome-wide CRISPR knockout sgRNAs were infected with HCoV-229E-mGreen (MOI 0.5, 24 h). Infected reporter-positive cells were sorted for genomic DNA extraction and subsequent sgRNA sequence analysis. Genes were analyzed using MAGeCK software and ranked based on -log10 (MAGeCK score) and P-values. **B.** Validation of the 12 top-ranked genes, with a cutoff of false discovery rate (FDR) < 0.05. A549 cells were edited with two independent sgRNAs per gene. Following infection with HCoV-229E (MOI 0.5, 24 h), infection efficiency was quantified by flow cytometry for the percentage of nucleocapsid (N)-positive cells. **C.** Representative images of control and *UHRF1*-knockout A549 cells infected with HCoV-229E (MOI 0.5, 24 h), and analyzed by Operetta high-content imaging. Scale bar, 100 μm. **D.** Cell viability of control and gene-knockout A549 cells was assessed 48 h after seeding. **E.** Overexpression of human UHRF1 inhibits HCoV-229E infection. HeLa cells overexpressing UHRF1 were infected with HCoV-229E (MOI 0.5, 24 h). Virus infection efficiency was determined by flow cytometry. **F.** Trans-complementation of UHRF1 in knockout cells inhibits HCoV-229E infection. A549 cells with or without exogenous UHRF1 expression were edited with sgRNA targeting endogenous UHRF1, and infected with HCoV-229E (MOI 0.5, 24 h). Error bars represent standard deviations from three independent experiments. Two-way ANOVA with Sidak’s test (B and F); unpaired t-test (D and E). \**P* < 0.05; \*\*\**P* < 0.001; \*\*\*\**P* < 0.001; ns, not significant.

We selected the top 12 genes for validation, using a cutoff of FDR < 0.05. For each gene target, A549 cells were transduced with two independent sgRNAs and subsequently infected with HCoV-229E. Notably, knockout of *UHRF1* resulted in a substantial increase in HCoV-229E infection, from approximately 1% to nearly 80%, compared to control cells **(Fig. 1B and C)**. The next candidate, *STARD3*, showed an increase in infectivity to over 20% **(Fig. 1B)**. STARD3 has been previously reported as a negative regulator for SARS-CoV-2 infection^23^. Thus, *UHRF1* emerged as our most compelling target, given its previously unknown role in coronavirus infection.

The observed enhancement of HCoV-229E infection was not due to cytotoxicity caused by *UHRF1* knockout **(Fig. 1D)**. As expected, overexpression of *UHRF1* cDNA in permissive HeLa cells led to a reduction in virus infection **(Fig. 1E)**. Furthermore, the restriction effect of UHRF1 in knockout cells could be rescued by genetic complementation **(Fig. 1F)**. These findings highlight the significant role of UHRF1 in restricting HCoV-229E infection.

### UHRF1 specifically restricts the infection of coronaviruses

To determine whether UHRF1 acts as a restriction factor across different coronavirus genera, we infected control and *UHRF1*-knockout cells with representatives from all four genera: the alpha genus including HCoV-NL63 and swine acute diarrhea syndrome coronavirus (SADS-CoV) **(Fig. 2A and Supplementary Fig. 1B)**, the beta genus SARS-CoV-2 and HCoV-OC43 **(Fig. 2B and Supplementary Fig. 1C)**, the gamma genus including infectious bronchitis virus (IBV) **(Fig. 2C)**, and the delta genus including porcine deltacoronavirus (PDCoV) **(Fig. 2D)**. Following *UHRF1* knockout, infection rates increased by 2- to 3-fold for most tested coronaviruses compared with control cells, except for HCoV-NL63 and SADS-CoV, whose infectivity was too low to assess precisely. However, it should be noted that the enhancement effect of *UHRF1* knockout was less pronounced for these coronaviruses compared to HCoV-229E **(Fig. 1)**. Additionally, trans-complementation of *UHRF1* cDNA in knockout cells restored the inhibitory effects on SARS-CoV-2 and PDCoV (**Supplementary Fig. 1D**).

**Figure 2.**
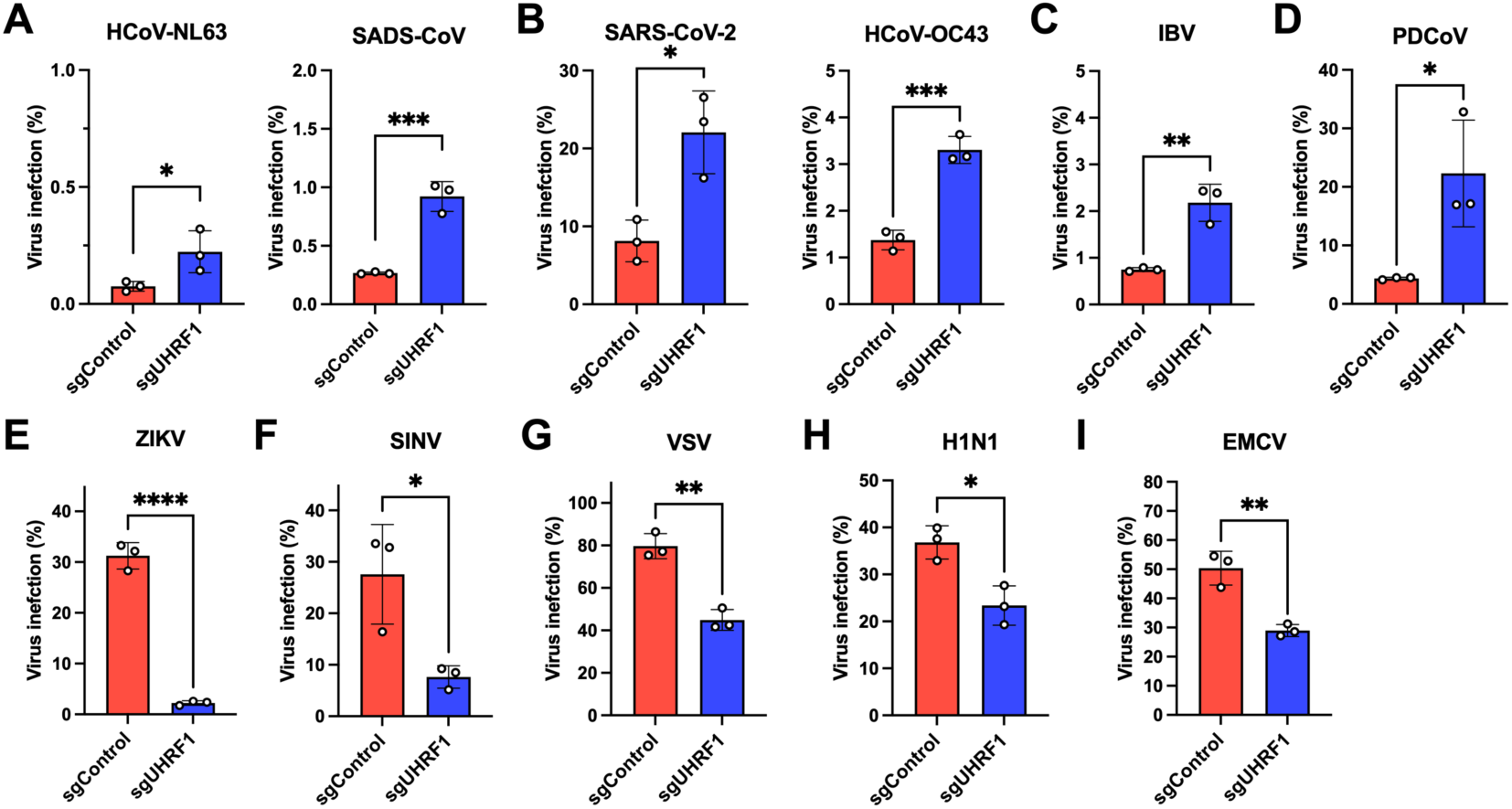
UHRF1 specifically restricts the infection of coronaviruses. **A-D.** Validation of UHRF1 as a restriction factor across coronavirus genera. Control and *UHRF1*-knockout A549-ACE2 cells were infected with alphacoronaviruses (HCoV-NL63, MOI 1, 24 h; SADS-CoV, MOI 1, 24 h) and betacoronavirus (SARS-CoV-2, MOI 0.1, 24 h). *UHRF1*-knockout HeLa cells were challenged with betacoronavirus (HCoV-OC43, MOI 1, 24 h), gammacoronavirus (IBV, MOI 1, 24 h), and deltacoronavirus (PDCoV, MOI 0.3, 24 h). Infection efficiency was analyzed by flow cytometry for the percentage of N-positive cells. **E-I.** Proviral effect of UHRF1 on unrelated RNA viruses. *UHRF1*-knockout A549 cells were infected with ZIKV (MOI 1, 24 h), SINV (MOI 3, 24 h), VSV (MOI 1, 15 h), H1N1 (MOI 1, 24 h), and EMCV (MOI 0.1, 10 h). Infection efficiency was determined by flow cytometry for the percentage of viral-positive cells. Error bars represent standard deviations from three independent experiments. Unpaired t-test; \**P* < 0.05; \*\**P* < 0.01; \*\*\**P* < 0.001; \*\*\*\**P* < 0.001; ns, not significant.

Previous studies have reported that UHRF1 deficiency negatively regulates antiviral innate immunity, thereby inhibiting infection of VSV and influenza virus^18^, which appears contradictory to our findings of enhanced coronavirus infection. To clarify this, we infected *UHRF1*-knockout A549 cells with a range of unrelated enveloped RNA viruses, including Zika virus (ZIKV), Sindbis virus (SINV), VSV, influenza virus (H1N1), and the non-enveloped RNA virus encephalomyocarditis virus (EMCV) **(Fig. 2E-I)**. Intriguingly, *UHRF* knockout reduced infection by these viruses, highlighting a distinct restrictive effect of UHRF1 on coronaviruses compared to other RNA viruses.

As a recognized negative regulator of innate immunity^18,19,24^, it remains unclear whether UHRF1 influences HCoV-229E infection through the interferon pathway. To address this, we knocked out *UHRF1* in *IPS-1-* and *STAT1*-deficient clonal cell lines and infected them with HCoV-229E. The results showed that *UHRF1* knockout still significantly enhanced viral infection **(Supplementary Fig. 2A and B)**. Furthermore, treatment of *UHRF1*-knockout cells with Ruxolitinib, an inhibitor of the JAK/STAT signaling pathway, did not reduce the heightened infection efficiency of HCoV-229E **(Supplementary Fig. 2C)**. Similar results were observed for SARS-CoV-2 in *IPS-1-* and *STAT1*-deficient cells in which editing of UHRF1 increased the infection **(Supplementary Fig. 2D)**. Interestingly, ZIKV and SINV infections were still inhibited in *UHRF1*-edited cells in the *IPS-1-* and *STAT1*-knockout backgrounds. These findings suggest that UHRF1 may restrict the infection by HCoV-229E and SARS-CoV-2 independently of the IFN signaling pathway.

### UHRF1 inhibits HCoV-229E entry by suppressing the APN expression

To investigate the mechanism by which UHRF1 restricts coronavirus infection, particularly HCoV-229E, we examined the entry stage, which is critical for cell susceptibility, using a VSV-based pseudovirus system expressing coronavirus spike proteins. Knockout of *UHRF1* markedly increased the infectivity of pseudovirus containing the HCoV-229E spike protein by over 30-fold, highlighting its critical role in blocking HCoV-229E entry **(Fig. 3A)**. *UHRF1* knockout also enhanced SARS-CoV-2 pseudovirus infection by approximately 1.6-fold **(Fig. 3B)**. In contrast, entry mediated by the VSV glycoprotein (VSV-G) was reduced in *UHRF1*-knockout cells, consistent with previous findings that UHRF1 promotes VSV infection **(Fig. 3C)**^18^. These results suggest that UHRF1 is critical for blocking the entry of HCoV-229E.

**Figure 3.**
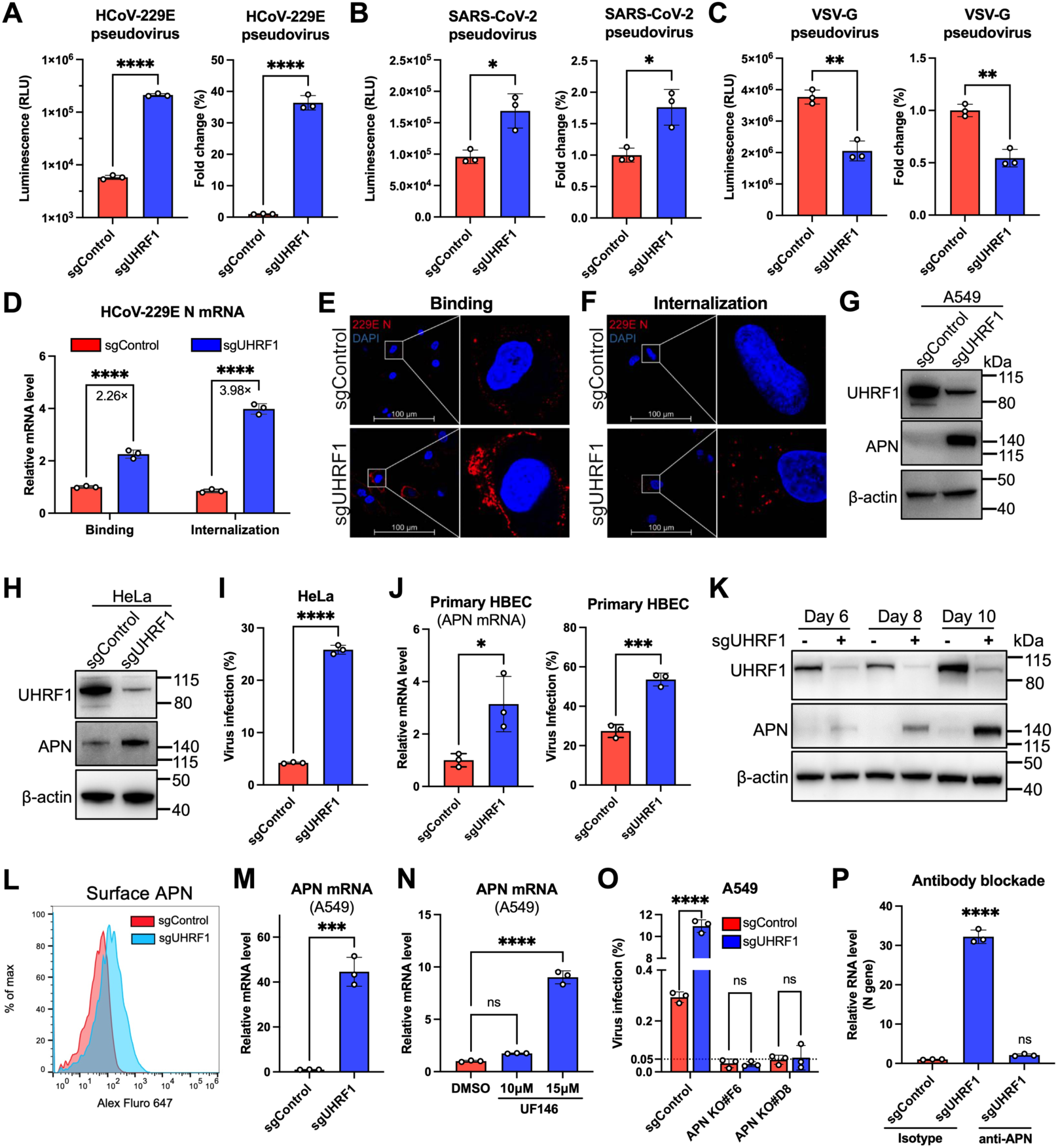
UHRF1 inhibits HCoV-229E entry by suppressing APN expression. **A-C.** Pseudovirus infection assays. Control and *UHRF1*-knockout A549-ACE2 cells were infected with VSV-based pseudoviruses bearing the spike protein of HCoV-229E (A), SARS-CoV-2 (B), or the glycoprotein of vesicular stomatitis virus (VSV-G) (C). Luciferase activity was measured at 12 h post-infection. **D-F.** Virus binding and internalization assays. Cells were incubated with HCoV-229E (MOI 10). Bound or internalized virions were quantified by qRT-PCR for genomic RNA or analyzed by confocal microscopy for N protein. **G-H.** Western blotting analysis of UHRF1 and APN expression in gene-knockout A549 or HeLa cells. β-actin was used as an internal control. **I.** Infection efficiency of HCoV-229E (MOI 0.5, 24 h) in control and *UHRF1*-knockout HeLa cells, determined by flow cytometry. **J.** Relative APN mRNA levels and infection efficiency of HCoV-229E (MOI 1, 12 h) in control and *UHRF1*-knockout primary human bronchial epithelial cells (HBEC). mRNA levels analyzed by qRT-PCR were normalized to internal control GAPDH and then to control cells. Infectivity was determined by flow cytometry. **K.** Temporal expression of APN in *UHRF1*-knockout A549 cells. Cell lysates were collected at different days post-transduction of sgRNA-expressing lentivirus and analyzed by western blotting for UHRF1, APN, and β-actin. **L.** Surface expression of APN analyzed by flow cytometry in A549 cells edited with control or *UHRF1* sgRNA. **M.** Relative APN mRNA levels analyzed by qRT-PCR in A549 cells edited with control or *UHRF1* sgRNA. mRNA levels were normalized to internal control GAPDH and then to control cells. **N.** Relative APN mRNA levels analyzed by qRT-PCR in A549 cells treated with UHRF1 inhibitor UF146 (10 or 15 μM) for 2 days. mRNA levels were normalized to internal control GAPDH and then to control cells. **O.** Effect of UHRF1 on HCoV-229E infection in *APN*-knockout A549 clonal cells. Control and two *APN*-knockout clonal cell lines were edited with control or *UHRF1* sgRNA, and infected with HCoV-229E (MOI 0.5, 24 h). The proportion of N-positive cells was detected by flow cytometry. **P.** Effect of APN-blocking antibody on HCoV-229E infection in *UHRF1*-knockout A549 cells. Gene-knockout cells were pre-treated with 5 μg/ml APN-blocking antibody or isotype control for 1 h, then infected with HCoV-229E (MOI 0.5, 24 h) in the presence of antibody. Total cellular RNA was extracted to determine relative levels of HCoV-229E N gene by qRT-PCR, and normalized to GAPDH. Error bars represent standard deviations from three independent experiments. Unpaired t-test (A-C, I-J, M); two-way ANOVA with Sidak’s test (D and O); one-way ANOVA with Sidak’s test (N and P). \**P* < 0.05; \*\**P* < 0.01; \*\*\**P* < 0.001; \*\*\*\**P* < 0.001; ns, not significant.

Next, we investigated how UHRF1 significantly impacts HCoV-229E entry. To determine whether UHRF1 restricts HCoV-229E binding to the cell surface, cells were incubated with the virus at 4℃ for 45 minutes and washed to remove unbound virions. qRT-PCR analysis revealed that HCoV-229E binding to *UHRF1*-knockout cells was 2.3-fold higher than in controls **(Fig. 3D)**. Confocal microscopy confirmed that more viral particles were bound on UHRF1-knockout cells **(Fig. 3E)**. To assess whether virion internalization was enhanced in *UHRF1*-knockout cells, cells were shifted to 37℃ for 45 minutes after binding, and treated with proteinase K to remove uninternalized viral particles before qRT-PCR analysis. Confocal microscopy was also used to evaluate virion internalization. The results showed that the amount of internalized viruses in *UHRF1*-knockout cells increased approximately 4-fold compared to controls **(Fig. 3D and F)**, indicating that UHRF1 restricts HCoV-229E entry at both the binding and internalization steps.

We then explored host factors regulated by UHRF1 that might inhibit HCoV-229E entry. Engagement with the primary receptor APN is critical for HCoV-229E entry into host cells. Notably, APN expression was undetectable by immunoblotting in control A549 cells but was significantly elevated in *UHRF1*-knockout cells **(Fig. 3G)**. Similar results were observed in HeLa cells, where APN was weakly detected in controls, and its upregulation correlated with increased susceptibility to HCoV-229E following *UHRF1* knockout **(Fig. 3H-I)**. Next, we edited the *UHRF1* in primary human bronchial epithelial cells (HBEC), resulting in increased expression of APN and enhanced HCoV-229E infection (**Fig.3J**). We also investigated the temporal regulation of APN expression by UHRF1. UHRF1 depletion was effective by day 6 post-transduction of CRISPR knockout sgRNA, but significant upregulation of APN expression was only observed by day 8 or 10 **(Fig. 3K)**, suggesting a delayed regulatory effect of UHRF1 on APN expression. Additionally, *UHRF1* knockout increased APN levels on the plasma membrane **(Fig. 3L)**. Knockout of *UHRF1* or treatment with UF146, a UHRF1 inhibitor that forms hydrogen bonds with the SRA domain groove of UHRF1^16^, significantly increased APN mRNA levels **(Fig. 3M and N)**.

To determine whether APN plays a key role in UHRF1-mediated regulation of HCoV-229E infection, we knocked out APN in A549 cells and selected clonal populations. Due to the challenge of detecting low endogenous APN expression, Inference of CRISPR Edits (ICE) analysis^25^ was used to assess knockout efficiency **(Supplementary Fig. 3)**. Two clones were selected for further experiments. Knockout of *UHRF1* in *APN*-knockout clonal cells completely abrogated the upregulatory effect of UHRF1 on HCoV-229E infection **(Fig. 3O)**. Additionally, *UHRF1*-knockout cells treated with an APN-blocking antibody before and during viral infection showed nearly complete loss of the enhanced effect of UHRF1 on HCoV-229E infection, as quantified by viral RNA levels using qRT-PCR **(Fig. 3P)**. These results demonstrate that UHRF1 restricts HCoV-229E infection primarily by downregulating APN receptor expression.

### UHRF1 inhibits HCoV-229E infection by maintaining APN promoter methylation

The UBL and RING domains of UHRF1 possess ubiquitinase and E3 ligase activities, respectively, and previous studies have shown that UHRF1 promotes the ubiquitination-mediated degradation of the tumor-suppressor protein promyelocytic leukemia protein^26^. To investigate how APN expression is regulated by UHRF1, we examined whether UHRF1 induces APN degradation. Transient expression of APN protein revealed that *UHRF1* knockout did not affect APN protein levels **(Fig. 4A)**.

**Figure 4.**
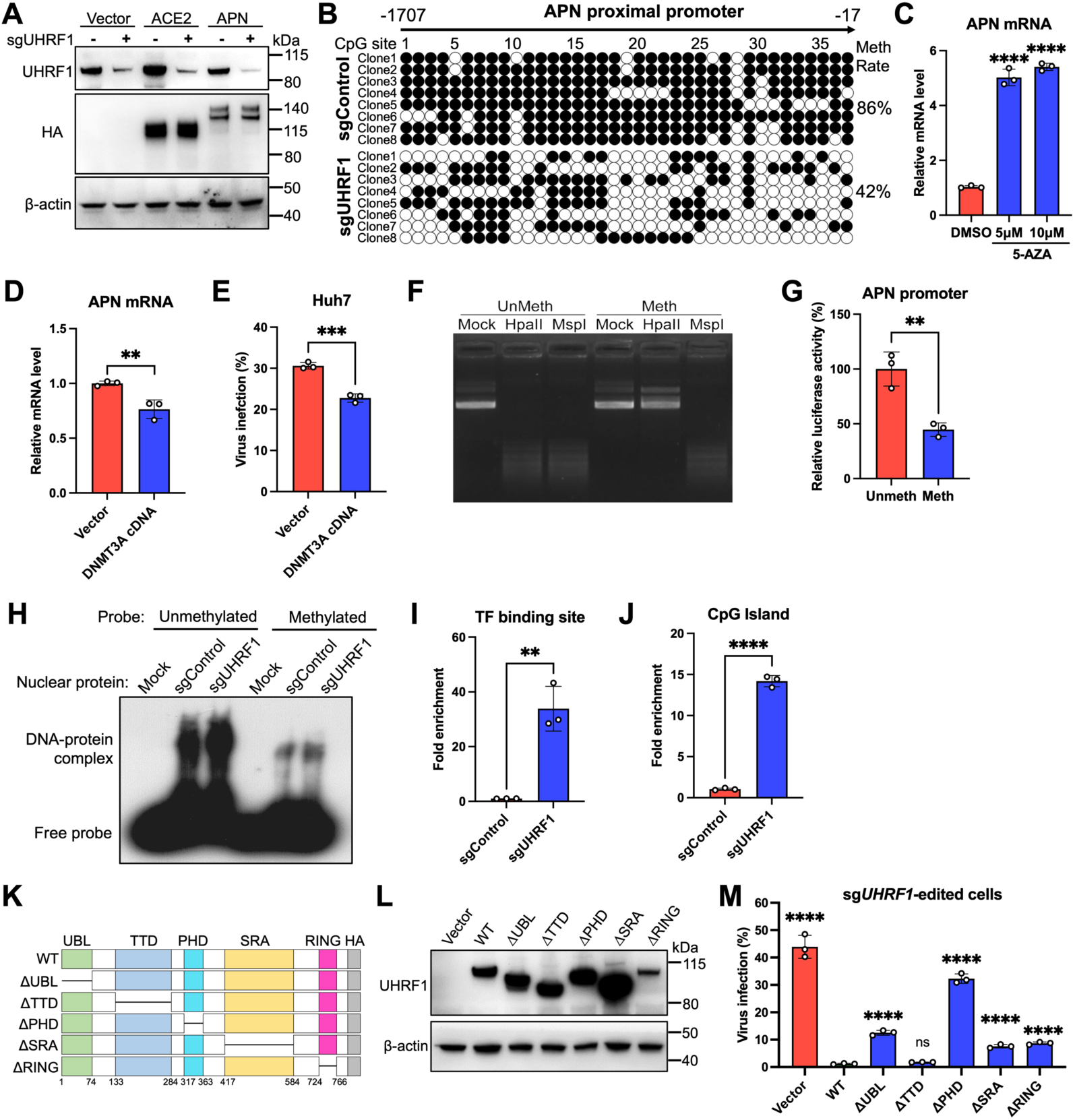
UHRF1 inhibits HCoV-229E infection by maintaining APN promoter methylation. **A.** UHRF1 does not regulate the expression of exogenous HA-tagged APN and ACE2. Control and *UHRF1*-knockout A549 cells were transiently transfected with plasmids expressing APN or ACE2, and cell lysates were analyzed by western blotting with anti-HA antibody. **B.** Bisulfite sequencing of CpG sites in APN proximal promoter from control and *UHRF1*-knockout A549 cells. The methylation (Meth) rate was calculated as the ratio of methylated sites to the total number of sites tested. **C.** Effects of 5-AZA treatment on APN mRNA levels. A549 cells were treated with 5 or 10 μM 5-AZA for 3 days, and total RNA was extracted to determine relative APN mRNA levels by qRT-PCR. GAPDH was used as an internal control. **D-E.** Effect of DNMT3A on APN expression and HCoV-229E infection. Huh7 cells stably expressing DNMT3A were generated by lentivirus transduction. Relative APN mRNA levels were determined by qRT-PCR (D), and infection efficiency was detected by flow cytometry after infection with HCoV-229E (MOI 0.01, 24 h) (E). **F-G.** *In vitro* methylation and dual-luciferase reporter assays. The luciferase reporter plasmid bearing APN promoter sequences was methylated by CpG methyltransferase M.SssI, and methylation status was verified by HpaII/MspI digestion (F). Unmethylated (Unmeth) or methylated (Meth) luciferase reporter plasmid was co-transfected with pRL-TK plasmid into HEK 293T cells, and luciferase activity was measured at 24 h post-transfection (G). Renilla luciferase was used as an internal control, and results were normalized to unmethylated plasmid. **H.** Electrophoretic mobility shift assay (EMSA). Nuclear extracts from control and *UHRF1*-knockout A549 cells were incubated with biotin-labeled unmethylated or methylated APN promoter probe to detect DNA-protein complexes. **I-J.** Chromatin immunoprecipitation (ChIP) assay with HA-tagged c-Maf expression in control and *UHRF1*-knockout A549 cells. qPCR was performed to detect c-Maf binding to the transcription factor (TF) binding site (I) or CpG island (J) of the APN proximal promoter. **K.** Schematic diagram of wild-type UHRF1 protein and its truncations with HA tag at the C terminus. **L-M.** Expression and function analysis of UHRF1 truncations in *UHRF1*-knockout A549 cells. Cell lines stably expressing wild-type or truncated UHRF1 were established by lentivirus transduction and verified by western blotting (L). Cells were infected with HCoV-229E (MOI 0.5, 24 h) at day 10 post-transduction, and the proportion of N-positive cells was determined by flow cytometry (M). Error bars represent standard deviations from three independent experiments. One-way ANOVA with Sidak’s test (C and M); unpaired t-test (D-E, G, I-J). \*\**P* < 0.01; \*\*\**P* < 0.001; \*\*\*\**P* < 0.001; ns, not significant.

Since UHRF1 has been reported to influence gene expression through promoter methylation^27,28^,we hypothesized that UHRF1 regulates APN expression at the transcriptional level by modulating promoter methylation. Using bisulfite cloning and sequencing, we assessed the methylation status of CpG sites in the APN promoter and observed a decrease in methylation rates from 86.15% to 42.23% **(Fig. 4B)**. This demethylation likely activates APN expression. Treatment of A549 cells with the DNA methylation inhibitor 5-azacytidine (5-AZA)^29^, siginficantly increased APN mRNA levels, further supporting the role of DNA methylation in regulating APN expression **(Fig. 4C)**.

To explore the impact of methylation on APN expression in a more susceptible cell line, we used Huh7 cells, which exhibit high APN expression^30^. Stable expression of DNMT3A, an enzyme that catalyzes the transfer of methyl groups to specific CpG structures in DNA^31^, in Huh7 cells decreased APN mRNA levels and reduced HCoV-229E infection compared to empty vector controls **(Fig. 4D and E)**. An *in vitro* methylation assay using a luciferase-based reporter vector containing the APN proximal promoter showed that the unmethylated promoter led to a significantly higher luciferase activity than the methylated promoter **(Fig. 4F and G)**, indicating direct regulation of APN expression by CpG methylation.

To further elucidate how DNA methylation suppresses APN transcription, we examined the binding of transcription factors to the methylated promoter. Methylation of the APN promoter reduced the formation of DNA-protein complexes with nuclear proteins, as shown by electrophoretic mobility shift assay (EMSA) **(Fig. 4H)**. Protooncogene c-Maf is reported as a transcription factor for APN expression in epithelial cells^32^. Chromatin immunoprecipitation (ChIP)-qPCR revealed that *UHRF1* knockout promotes the recruitment of c-Maf to the APN promoter **(Fig. 4I and J)**. However, no differences were observed in the enrichment acetylated histones (H3K9ac, H3K14ac, and H3K27ac) on the APN promoter between control and *UHRF1*-knockout cells **(Supplementary Fig. 4A and B)**, suggesting that promoter methylation, rather than histone modifications, inhibits APN transcription.

To identify the critical structural domains of UHRF1 responsible for restricting HCoV-229E infection, we constructed HA-tagged truncations of UHRF1^20^, and confirmed their expression by immunoblotting **(Fig. 4K and L)**. Complementation assays showed that all domains except the TTD domain were indispensable for regulating HCoV-229E infection **(Fig. 4M)**. Previous studies have shown that PHD and SRA domains of UHRF1 are required to maintain DNA hypermethylation^33^, and both UBL and RING domains are also critical for proper nuclear localization of DNMT1 and maintenance of DNA methylation ^34^. But global DNA methylation is largely independent of H3K9 methylation with TTD mutation^35^. Collectively, these findings support the conclusion that UHRF1 inhibits HCoV-229E infection by maintaining APN promoter methylation.

### APN is the most dominant gene suppressed by UHRF1 for HCoV-229E infection

To comprehensively elucidate the role of UHRF1-regulated genes in coronavirus infection, we performed RNA-seq analysis of control and *UHRF1*-knockout cells. Using a cuttoff of log_2_ fold change >2 and P-value <0.05, we identified 2210 upregulated and 51 downregulated genes in knockout cells compared to controls **(Fig. 5A)**. APN mRNA was upregulated approximately 40-fold in *UHRF1*-knockout cells. Using the STRING database, we constructed an interaction network of UHRF1-regulated genes and extracted the top 20 hub genes using the Cytohubba plugin^36,37^. These genes are primarily involved in pathways related to inflammation, innate immunity, and metabolic processing **(Supplementary Fig. 5A)**, consistent with previous reports that UHRF1 acts as a negative regulator of innate immune signalling^24^. However, these pathways do not explain the enhanced coronavirus infection observed in *UHRF1*-knockout cells. KEGG and GO enrichment analysis also did not identify pathways that benefit viral infection **(Supplementary Fig. 5B and C)**.

**Figure 5.**
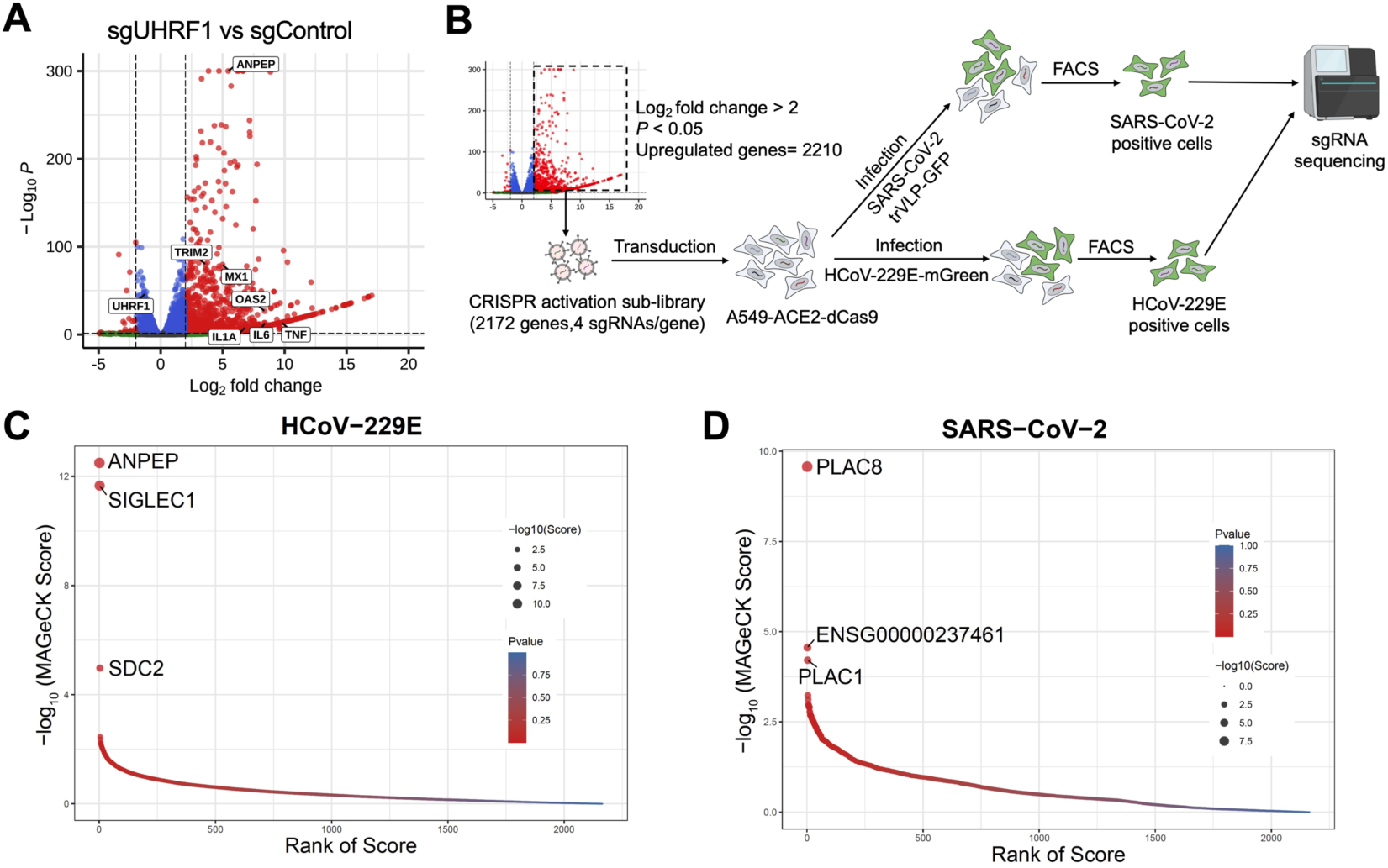
APN is the most dominant gene suppressed by UHRF1 for HCoV-229E infection. **A.** Volcano plot of RNA-seq analysis. Total cellular RNA was extracted from control and *UHRF1*-knockout A549-ACE2 cells and subjected to RNA-seq. Genes with an absolute Log_2_ fold change >2 and adjusted P-value <0.05 were considered as differentially expressed. **B.** Schematic of focused CRISPR activation screening. A sub-library targeting 2172 of the 2210 upregulated genes identified from RNA-seq analysis of *UHRF1*-knockout cells, with ∼4 sgRNAs per gene, were generated and transduced into A549-ACE2-dCas9 cells. Cells were infected with HCoV-229E-mGreen (MOI 0.5, 24 h) or SARS-CoV-2 transcription- and replication-competent virus-like particles in which the N gene is replaced by the reporter GFP (trVLP-GFP)^38^ (MOI 0.5, 24 h). Infected reporter-positive cells were sorted for genomic DNA extraction and sgRNA sequence analysis. **C-D.** Genes identified from CRISPR screens for HCoV-229E (C) and SARS-CoV-2 (D). Genes were analyzed by MAGeCK software and sorted based on -log_10_ (MAGeCK score) and P-values.

To systemically identify the UHRF1-suppressed genes that promote coronavirus infection, we constructed a focused CRISPR activation sub-library containing 2172 of the 2210 upregulated genes identified in the RNAseq analysis of *UHRF1*-knockout cells **(Fig.5B)**. The A549 cell library were infected with HCoV-229E-mGreen virus, and reporter-postive cells were sorted to for data analysis **(Fig. 5B)**. As a control, the library was also infected with transcription- and replication-competent SARS-CoV-2 virus-like particles (SARS-CoV-2 trVLP-GFP), in which the nucleocapsid (N) gene is replaced by the GFP reporter gene and the particles are trans-packaged in N-expressing cells^38^. Intriguingly, APN scored the highest in the HCoV-229E screen, followed by Sialic Acid Binding Ig Like Lectin 1 (*SIGLEC1*) and Syndecan 2 (SDC2), which have been previously reported to favor coronavirus infection^39,40^ **(Fig. 5C)**. For SARS-CoV-2, the top-ranked gene was PLAC8, followed by ENSG00000237461 and PLAC1 **(Fig. 5D)**. PLAC8 has been identified as a proviral factor for SADS-CoV and SARS-CoV-2 infection^41,42^. Despite their similar names, PLAC1 and PLAC8 have distinct 3D structures, as predicted by AlphaFold 2, suggesting possibly divergent proviral mechanisms **(Supplementary Fig. 5D)**. ENSG00000237461, also known as LOC101928438, is a gene from the lncRNA family with unknown function. These findings confirm that APN is the most prominent gene suppressed by UHRF1 for HCoV-229E infection, and additional proviral genes, like SIGLEC1, SDC2, and PLAC8, are also regulated.

We further investigated the influence of UHRF1 on other common entry factors of coronaviruses. No changes were observed in the expression of CTSL, TMPRSS2, Furin, ACE2, DC-SIGN, or TIM-1 in knockout cells **(Supplementary Fig. 6A and B)**. However, we observed increased surface expression of heparan sulfate (HS), a known common adhesion factor for coronaviruses^43^ **(Supplementary Fig. 6C)**. Additionally, increased levels of heparan sulfate were detected in the supernatants and cell lysates **(Supplementary Fig. 6D)**. qRT-PCR analysis showed that genes involved in HS biosynthesis, such as XYLT1,EXT1 and NDST1, are upregulated in *UHRF1*-knockout cells compared to controls **(Supplementary Fig. 6E)**. Furthermore, increased SARS-CoV-2 infection by *UHRF1* knockout was attenuted in *B3GAT3*-deficient cells (**Supplementary Fig. 6F**). These results suggest that UHRF1 also regulats other common entry factors, e.g., heparan sulfate, albeit with a less pronouced effect, acting as a coronavirus restrictor.

### Age-related susceptibility to HCoV-229E driven by UHRF1 epigenetic modulation

To determine whether UHRF1 correlates with age-related susceptibility to HCoV-229E, we analyzed transcriptomic data of lung tissue in the GTEx database. The results indicated that UHRF1 expression decreases with age, with mean normalized transcripts per million (nTPM) approximately 50% lower in individuals aged 60-79 years compared to those aged 20-39 years **(Fig. 6A)**. Conversely, APN expression increased with age, with mean nTPM rising from 30.52 in 20-39 year-olds to 34.98 in 60-79 year-olds **(Fig. 6B)**. Correlation analysis using the ggstatsplot R package revealed a negative correlation between UHRF1 expression and age, and a positive correlation between APN expression and age **(Fig. 6C)**.

**Figure 6.**
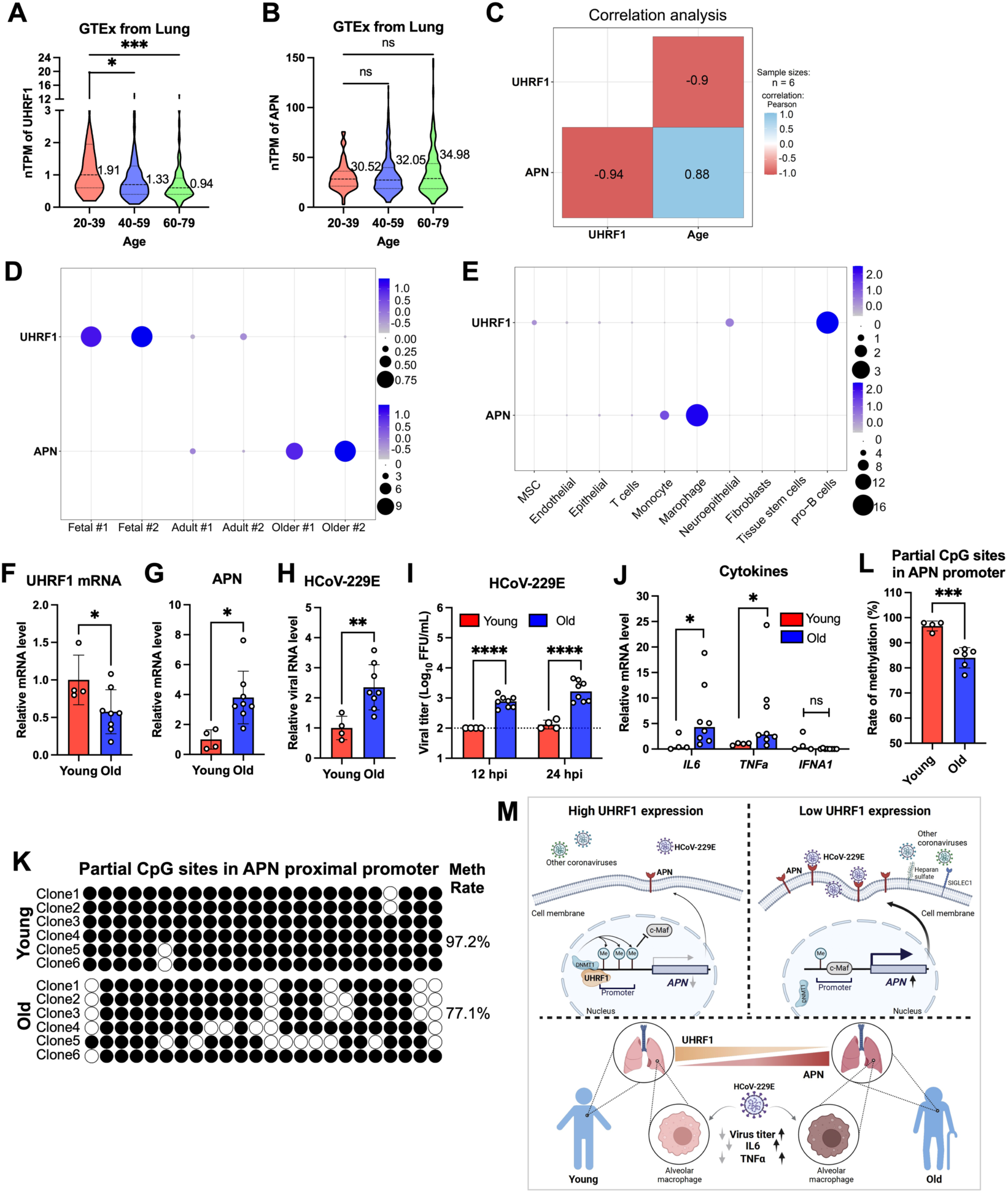
Age-related susceptibility to HCoV-229E driven by UHRF1 epigenetic modulation. **A-B.** Transcriptomic analysis of UHRF1 (A) and APN (B) expression in lung tissues from three age groups. **C.** Correlation analysis of UHRF1 and APN transcription levels with age. **D.** UHRF1 and APN expression levels in single-cell RNA-seq (scRNA-seq) data of lung tissues from three age groups. Dot plot shows the percentage of cells in each population with detectable gene expression. Dot size indicates the percentage of positive cells, and color indicates mean expression levels. **E.** Analysis of UHRF1 and APN gene expression in scRNA-seq data of various lung cell types. Dot size indicates the percentage of positive cells, and color indicates the mean expression levels. **F-G.** Expression levels of UHRF1 (F) and APN (G) in alveolar macrophages from young (n=4) and old (n=8) donors. Relative mRNA levels were determined by qRT-PCR, and expressed relative to young donors. **H-J.** HCoV-229E infection in alveolar macrophages from young and old donors. At 24 h post infection, viral RNA in cells was determined by qRT-PCR (H); At 12 or 24 h post infection (hpi), virus production in the supernatant was titrated by focus-forming assay (I). Cytokine expression in cells was determined by qRT-PCR (J); **K-L.** Bisulfite sequencing of CpG sites in APN proximal promoter from one representative (K) or pooled (L) young and old donors. Methylation (Meth) rates were calculated as the ratio of methylated sites to the total number of sites tested. **M.** Schematic of the role of *UHRF1* in coronavirus infection. UHRF1 maintains DNA methylation by recruiting the methyltransferase DNMT1 to gene promoters, thereby regulating gene expression. Hypermethylation of the APN promoter prevents transcription factors, such as c-Maf, from initiating transcription, leading to downregulation of APN protein expression. Since APN serves as the key entry receptor for HCoV-229E, this suppression restricts viral infection. Conversely, hypomethylation of the APN promoter upregulates APN expression, promoting HCoV-229E infection. Additionally, UHRF1 regulates other proviral host factors, including SIGLEC1, PLAC8, and heparan sulfate biosynthesis genes, which influence coronavirus infection. UHRF1 expression decreases with age, negatively correlating with increased APN levels. This age-related decline in UHRF1 was observed in primary alveolar macrophages isolated from elderly individuals, which exhibited heightened susceptibility to HCoV-229E infection and increased pro-inflammatory cytokine production compared to those from younger individuals. Thus, UHRF1 restricts coronavirus infection through epigenetic silencing of host factors, particularly the entry receptor APN for HCoV-229E. Error bars represent standard deviations (F-I); the median values are shown (I, J). One-way ANOVA with dunnett’s test (A-B); unpaired t test (F-I, L); Mann-Whitney test (J). \**P* < 0.05; \*\**P* < 0.01; \*\*\**P* < 0.001; \*\*\*\**P* < 0.001; ns, not significant.

Further analysis of single-cell RNA sequencing (scRNA-seq) data across different age groups showed that UHRF1 is highly expressed in fetal lungs but exhibits reduced expression in older individuals. In contrast, APN is highly expressed in the lungs of older individuals, possibly contributing to their increased susceptibility to HCoV-229E **(Fig. 6D)**. Among various lung cell types, APN was predominantly expressed in macrophages and monocytes **(Fig. 6E)**, consistent with prior findings that these cells are targets for HCoV-229E infection^7,8^. These results underscore a negative correlation between UHRF1 expression and age, which may explain the heightened sensitivity of the elderly to HCoV-229E infection.

Macrophages are pivotal in regulating viral infections and inflammation^10,11^, serving as target cells for HCoV-229E. To investigate the increased susceptibility and severity of HCoV-229E infection in older individuals, we isolated primary alveolar macrophages from bronchoalveolar lavage fluid of young (20-25 years) and old (53-80 years) individuals with noninflammatory or noninfectious conditions. Consistent with the transcriptomic data, UHRF1 mRNA levels were reduced by 50% in the old group **(Fig. 6F)**, while APN mRNA levels, negatively regulated by UHRF1, increased approximately 4-fold **(Fig. 6G)**. Upon HCoV-229E infection, alveolar macrophages from elderly individuals exhibited significantly higher viral genome copy numbers in cells and titers in the supernatants compared to the young individuals **(Fig. 6H and I)**. Additionally, a significant increase in cytokines, such as *IL6* and *TNFα*, was observed in the old group **(Fig. 6J and Supplementary Fig. 6G)**. We also examined the methylation status of the APN promoter in alveolar macrophages and found reduced methylation in old individuals compared to the young group **(Fig. 6K and L)**. These results suggest that loss of UHRF1-regulated APN promoter methylation leads to increased APN expression in alveolar macrophages, promoting cell susceptibility to HCoV-229E infection and increasing the risk of severe disease in the elderly **(Supplementary Fig. 7)**.

## DISCUSSION

The emergence of SARS-CoV-2 has highlighted the devastating threat posed by coronaviruses to public health, particularly for the elderly population. Similarly, common human coronaviruses, such as HCoV-229E, can cause severe diseases in older individuals. To better understand viral-host interactions and identify host factors associated with viral pathogenesis and disease progression, in this study, we performed a genome-wide CRISPR knockout screen using HCoV-229E as a model virus. This screen revealed UHRF1 as a potent host restriction factor that inhibits HCoV-229E infection by suppressing the expression of its cell entry receptor APN.

The role of UHRF1 in viral infections has been previously reported. For instance, UHRF1 promotes the maintenance of HIV-1 latency by controlling Tat-mediated transcriptional activation and binding to the HIV-1 promoter for methylation^20,21^. Together with DNA methyltransferases DNMT1 and DNMT3B, UHRF1 maintains EBV latency by restricting oncoprotein expression^22^. Additionally, UHRF1 deficiency inhibits infection by alphaherpesviruses, influenza virus, and VSV, acting as a negative regulator of innate immunity^18,19^. Indeed, in our study, we found that UHRF1 is a proviral factor for some RNA viruses, including ZIKV, SINV, VSV, influenza virus, and EMCV.

However, we primarily identified UHRF1 as a potent host restriction factor for HCoV-229E infection, a role that extends to other coronaviruses. Although *UHRF1* knockout enhances host innate immunity, which can inhibit coronavirus infection, coronaviruses encode numerous genes that antagonize innate immune responses^44–46^. Importantly, the entry receptor APN^12,47^, which plays a key role in cell susceptibility to HCoV-229E, was markedly upregulated in *UHRF1*-knockout cells. To further elucidate UHRF1-regulated proviral factors, we performed a focused CRISPR activation screen of UHRF1-suppressed genes and confirmed the prominent role of the APN receptor in HCoV-229E infetion. Addtionally, other genes with pan-coronaivrus proviral functions, like SIGLEC1, SDC2, and PLAC8, were identified^39–42^. Similary, the expression of heparan sulfate, a broad-spectrum viral adhesion factor^48–52^, was significantly increased in knockout cells, potentially contributing to the restriction function of UHRF1 for multiple coronaviruses.

Hypomethylation of the APN promoter resulting from UHRF1 depletion activates APN expression. Although SIGLEC1, SDC2, PLAC8, and other genes involved in heparan sulfate biosynthesis are also regulated by UHRF1, their impact on infection by other coronaviruses is relatively weak compared to HCoV-229E. This variability can be attributed to the key host factors reguated by UHRF1 for specific viruses. Additionally, DNA methylation affects different genes at varying levels, as gene expression is regulated by a complex network of DNA methylation and histone modifications^53–55^. While mapping the domains of UHRF1 that regulate HCoV-229E infection, we found that all domains except the TTD domain are essential. The role of the TTD domain in maintaining DNA methylation is controversial ^33^, and our results suggest that it is not required for UHRF1 to restrict HCoV-229E infection.

Analysis of transcriptomic and scRNA-seq data indicated that UHRF1 expression decreases with age and negatively correlates with APN expression. Elevated APN expression due to UHRF1 downregulation may render the cells, particularly target cells such as (alveolar) macrophages and monocytes, more susceptible to HCoV-229E infection, leading to higher viral loads in the lungs of older individuals. Increased infection is potentially associated with elevated inflammation levels, even triggering inflammatory storms and rapidly progressing to acute respiratory distress syndrome^5^. To corroborate this, we isolated alveolar macrophages from young and elderly individuals and confirmed decreased UHRF1 and elevated APN expression in the elderly group. Consistent with this, HCoV-229E infection resulted in significantly higher viral titers in alveolar macrophages from elderly individuals compared to the young group, accompanied with increased cytokine production. These findings underscore that UHRF1 is an important age-related coronavirus host restriction factor.

Numerous genome-scale CRISPR screens have been performed to identify proviral or antiviral host factors for coronaviruses, particularly the SARS-CoV-2^23,56–61^. To our knowledge, no prior CRISPR screens have targeted antiviral host factors for HCoV-229E. In this study, our use of HCoV-229E as a model virus in a genome-wide CRISPR knockout screen, combined with focused activation screens, revealed UHRF1 as an age-related coronavirus restriction factor primarily through the regulation of receptor APN expression. The identification of UHRF1 highlights the host intrinsic defense systems against coronavirus infection via epigenetic regulation of key entry receptors and provides a new perspective for developing host-directed therapeutic strategies.

## METHODS

### Cells and viruses

A549 (ATCC #CCL-185), A549-ACE2^62^, HeLa (ATCC #CCL-2), Vero E6 (Cell Bank of the Chinese Academy of Sciences, Shanghai, China), HEK 293T (ATCC #CRL-3216), Huh7, swine testicular (ST), LLC-MK2, and HRT-18 cells were cultured at 37°C in Dulbecco’s Modified Eagle Medium (Hyclone #SH30243.01) supplemented with 10% fetal bovine serum (FBS), 10 mM HEPES, 1 mM Sodium pyruvate, 1× non-essential amino acids, and 100 U/ml of Penicillin-Streptomycin. The *APN*-knockout A549 clonal cell lines were generated by transducing packaged lentiCRISPR v2GFP (Addgene #82416) expressing representative sgRNA.The *IPS-* and *STAT1-*knockout A549 clonal cell lines were generated by transducing packaged lentiCRISPR v2 (Addgene #52961) or lentiCRISPR v2 Blast (Addgene #98293), respectively. Clonal cell lines were obtained by limiting dilution and verified by western blotting or Inference of CRISPR Edits (ICE) analysis^25^. All cell lines were tested routinely and free of mycoplasma contamination. HCoV-229E was propagated in Huh7 cells and titrated by focus-forming assay in Huh7 cells. HCoV-NL63 (LLC-MK2), swine acute diarrhea syndrome coronavirus (SADS-CoV) (Vero E6), SARS-CoV-2 (nCoV-SH01-Sfull) (Vero E6), HCoV-OC43 (HRT-18), infectious bronchitis virus (IBV) (Vero E6), porcine deltacoronavirus (PDCoV) (ST), Zika virus (ZIKV) (Vero E6), Sindbis virus (SINV) (BHK-21), vesicular stomatitis virus (VSV) (BHK-21), H1N1 influenza virus (MDCK), and encephalomyocarditis virus (EMCV) (HeLa) were prepared and titrated similarly in their respective cell lines, as indicated in parentheses. All experiments involving live SARS-CoV-2 virus were performed in a biosafety level 3 (BSL-3) facility of Fudan University.

### Genome-wide CRISPR knocout screen

The genome-wide CRISPR knockout library was generated as described previously^62^. Briefly, A549 cells expressing the Cas9 (A549-Cas9) were transduced with a packaged sgRNA lentivirus library targeting 19,114 genes (Addgene #73178)^63^, at a multiplicity of infection (MOI) of ∼0.3 by spinoculation at 1000 x g and 32°C for 30 min in 12-well plates. After selection with puromycin for around 7 days, cells were inoculated with HCoV-229E-mGreen, in which the mGreenLantern (mGreen) reporter gene replaces the ns4a. After infection at an MOI of 0.5 for 24 h, cells were harvested, and infected reporter-positive populations were sorted by flow cytometry. Genomic DNA from both sorted and uninfected cells was extracted for sgRNA amplification and next generation sequencing using an Illumina NovaSeq 6000 platform. sgRNA sequences were trimmed using the FASTX-Toolkit and cutadapt 1.8.1, and sgRNA abundance and gene ranking were analyzed using the MAGeCK computational tool **(see Supplementary Table 1)**.

### Gene validation

The top 12 genes with a false discovery rate (FDR) of less than 0.05 from the MAGeCK analysis were selected for validation. Two independent sgRNAs per gene were chosen from the CRISPR knockout library and cloned into the lentiCRISPR v2 (Addgene #52961). Lentiviruses were packaged with psPAX2 (Addgene #12260) and pMD2.G (Addgene #12259). A549 cells were transduced with lentiviruses expressing individual sgRNAs and selected with puromycin for 7 days. Gene-knockout cells were infected with HCoV-229E (MOI 0.5, 24 h), fixed, and stained for nucleocapsid (N) protein, followed by high-content imaging or flow cytometry. sgRNA sequences used for validation are listed in **Supplementary Table 2.**

Other coronaviruses and unrelated RNA viruses were selected to examine the antiviral specificity of UHRF1. Control and *UHRF1*-knockout A549-ACE2 cells were infected with alphacoronaviruses (HCoV-NL63, MOI 1, 24 h; SADS-CoV, MOI 1, 24 h) and betacoronavirus (SARS-CoV-2, MOI 0.1, 24 h). The validation was also performed in UHRF1-knockout A549 cells without ectopic expression of ACE2 for HCoV-NL63 (MOI 1, 24 h) and SARS-CoV-2 transcription- and replication-competent virus-like particles^64^, in which the N gene is replaced by reporter NanoLuc luciferase (trVLP-Nluc) (MOI 0.5, 24 h). UHRF1-knockout HeLa cells were challenged with betacoronavirus (HCoV-OC43, MOI 1, 24 h), gammacoronavirus (IBV, MOI 1, 24 h), and deltacoronavirus (PDCoV, MOI 0.3, 24 h). For unrelated RNA viruses, *UHRF1*-knockout A549 cells were infected with ZIKV (MOI 1, 24 h), SINV (MOI 3, 24 h), VSV (MOI 1, 15 h), H1N1 (MOI 1, 24 h), and EMCV (MOI 0.1, 10 h).

For validation in primary human bronchial epithelial cells (HBEC) (Procell #CP-H009)^65,66^, cells were prepared from fresh human bronchi of healthy donors. Human bronchi were dissected into 1- to 2-mm segments and digested with collagenase. Dissociated cells were pelted, washed, and cultured at 37°C with 5% CO2. The identity of isolated HBECs was confirmed by cell-surface staining of cytokeratin 19 expression. HBEC were transduced with lentiviruses expressing control or UHRF1-specific sgRNA and selected with puromycin for 6 days. Gene-edited cells were infected with HCoV-229E (MOI 1, 12 h), fixed, and stained for nucleocapsid (N) protein, followed by flow cytometry. Total RNAs were also extracted using RNAsimple Total RNA Kit (Tiangen #DP430) according to the manufacturer’s instructions. Relative expression of APN mRNA was analyzed by qRT-PCR as described below.

### Cell viability assay

Cell viability was assessed using the CellTiter-Lumi™ Steady Cell Viability Assay kit (Beyotime # C0069M) according to the manufacturer’s instructions. Approximately 1×10^4^ control and *UHRF1*-knockout A549 cells were seeded into opaque-walled 96-well plates. After 48 hours, CellTiter-Lumi™ reagent was added to each well, followed by shaking for 2 minutes. After a 10-minute incubation at room temperature, luminescence was measured using a FlexStation 3 (Molecular Devices) with an integration time of 0.5 seconds per well.

### Pseudotyped virus experiment

Vesicular stomatitis virus (VSV)-based pseudoviruses were produced in HEK 293T cells. Cells were transfected with a pcDNA3.1 vector expressing the full spike gene of HCoV-229E, the spike of SARS-CoV-2 lacking the C-terminal 21 amino acids, or VSV-G (pMD2.G, Addgene #12259), using Fugene^®^HD transfection reagent (Promega) for 24 hours. Cells were infected with single-cycle scVSV^ι1G^-Nluc-GFP^67^, in which the glycoprotein gene is deleted, at an MOI of 1 for 2 hours. After 3 washes, cells were maintained in culture medium with anti–VSV-G neutralizing antibody for 24 h. Supernatants were collected, centrifuged at 3,500 rpm at 4°C for 15 min to remove cell debris, and then aliquoted for storage at −80°C. Virus entry was assessed by transducing 30 µl of pseudoviruses into control and *UHRF1*-knockout A549-ACE2 cells in 96-well plates. After 12 hours, luciferase activity was measured using the Nano-Glo® Luciferase Assay Kit (Promega #N1120), and luminescence was recorded using a FlexStation 3 (Molecular Devices).

### Virus binding and internalization assays

To assess viral binding and internalization, control and *UHRF1*-knockout A549 cells were collected using TrypLE (Thermo #12605010) and pre-chilled on ice for 10 minutes before incubation with ice-cold HCoV-229E (MOI 10) for 45 minutes. Cells were washed three times with ice-cold PBS, lysed in Buffer RL (Tiangen #DP430) for RNA extraction, and analyzed by qRT-PCR. For internalization, after initial virus binding, cells were incubated at 37°C for 45 minutes. Uninternalized virions were removed by treating with 400 μg/ml protease K on ice for 45 minutes, followed by three PBS washes, cell lysis, and qRT-PCR analysis. The relative amount of bound and internalized virions was normalized to the internal control GAPDH.

For confocal microscopy, cells seeded on coverslips were incubated with HCoV-229E (MOI 10) on ice for 45 minutes. After three PBS washes, cells were fixed with 2% paraformaldehyde for 10 minutes, blocked with PBS containing 5% BSA and 0.3 M glycine for 2 hours at room temperature, then incubated overnight at 4°C with home-made mouse anti-HCoV-229E nucleocapsid (N) protein serum (1:1000) in PBS with 1% BSA. Cells were washed and incubated with goat anti-mouse IgG (H+L) conjugated with Alexa Fluor 555 (Thermo #A-21424, 2 μg/ml) for 2 hours at room temperature. For internalized virions, cells were shifted to 37°C for 45 minutes after binding, fixed, and stained in the presence of 0.2% saponin. Cells were counterstained with 4’,6-diamidino-2-phenylindole (DAPI), and images were captured using a Leica TCS SP8 confocal microscope and processed using Leica Application Suite X (LAS X, v3.7.0.20979) and ImageJ v2.0.0 (http://rsb.info.nih.gov/ij/).

### Surface staining

Control and *UHRF1*-knockout A549 cells were detached with TrypLE and incubated with primary antibodies against APN (Invitrogen #14-0138-82, 1 μg/ml), heparan sulfate (10E4) (USBiological #H1890, 1 μg/ml), ACE2 (Sino Biological #10108-RP01, 1:250), DC-SIGN (Biolegend #330102, 1 μg/ml), or TIM-1 (Biolegend #354002, 1 μg/ml) at 4°C for 25 minutes. After washing, cells were stained with goat anti-mouse or rabbit IgG (H+L) conjugated with Alexa Fluor 647 (Thermo Fisher #A21245, 2 μg/ml) for 25 minutes at 4 °C and analyzed by flow cytometry.

### Antibody blockade

Control and *UHRF1*-knockout A549 cells were seeded in 96-well plates. Following media removal, cells were treated with 5 μg/ml APN-blocking antibody (Invitrogen #14-0138-82) or isotype control for 1 hour, followed by infection with HCoV-229E (MOI 0.5, 24 h) in the presence of the antibody. Total cellular RNA was extracted, and viral RNA levels were quantified using qRT-PCR targeting the N gene of HCoV-229E.

### Bisulfite cloning & sequencing

DNA was extracted from control and *UHRF1*-knockout A549 cells using the DNeasy Blood & Tissue Kit (Qiagen #69506) and subjected to bisulfite conversion using the DNA Bisulfite Conversion Kit (Tiangen #DP215). Target fragments were amplified with the Methylation-specific PCR (MSP) Kit (Tiangen #EM101). PCR amplicons were cloned into the pCE2 TA/Blunt-Zero vector (Vazyme #C601-02), and eight colonies were selected for sequencing of CpG sites. DNA sequences of the APN proximal promoter underwent in silico bisulfite conversion using MethPrimer (http://www.urogene.org/cgi-bin/methprimer2/MethSequence.cgi). Primers used for PCR are listed in **Supplementary Table 2**.

### Chromatin immunoprecipitation (ChIP) assay

ChIP was performed using the ChIP Assay Kit (Beyotime #P2078) according to the manufacturer’s instructions. Briefly, control and *UHRF1*-knockout A549 cells were cross-linked with 1% formaldehyde for 10 min at 37°C, and the reaction was stopped by glycine solution for 5 min at room temperature. Cells were washed twice with cold PBS and harvested in SDS Lysis buffer containing 1mM PMSF. Samples were sonicated at 4°C (20% of max power, 30 s on and 30 s off for 3 min). Protein A+G Agarose/Salmon Sperm DNA was used to pre-clear the whole cell lysate for 30 min at 4°C. After 2% of the samples were extracted as an input control, the remaining samples were divided equally and incubated with anti-Histone H3 (acetyl K9) (Abcam #ab32129), anti-Histone H3 (acetyl K27) (Abcam #ab177178), and anti-Histone H3 (acetyl K14) (Abcam #ab52946) at 4°C overnight. To assess the transcription factor c-Maf, a similar procedure was performed with an anti-HA tag antibody (Proteintech #51064-2-AP). Protein A+G Agarose/Salmon Sperm DNA was added and incubated at 4°C for 60 min. Beads were washed sequentially with Low-Salt Immune Complex Wash Buffer, High-Salt Immune Complex Wash Buffer, LiCl Immune Complex Wash Buffer, and TE Buffer (twice) for 5 min at 4°C with rotation. DNA-protein complexes were eluted with elution buffer (1% SDS and 0.1 M NaHCO3), and de-crosslinked by adding 0.2 M NaCl and heating at 65°C for 4 hours. Proteins were digested with 40 μg/ml proteinase K, 10 mM EDTA, and 40 mM Tris (pH 6.5) for 1 hour at 45 °C. DNA segments were purified and used for qPCR. Primers are listed in **Supplementary Table 2**.

### Electrophoretic mobility shift assay (EMSA)

EMSA was performed as previously described^68^. APN promoter region (-207 to -20) were amplified from the genomic DNA of A549 cells by PCR and methylated using CpG Methyltransferase M.SssI (NEB #M0226S). Unmethylation or methylation probes were biotinylated using the Biotin 3’ End Labeling Kit (Beyotime #GS008). A Chemiluminescent EMSA kit (Beyotime #GS009) was used to detect interactions between the probes and nuclear proteins extracted from control and *UHRF1*-knockout A549 cells using NE-PER™ Nuclear and Cytoplasmic Extraction Reagents (Thermo #78833).

### *In vitro* methylation and dual-luciferase reporter assays

The APN promoter sequences (-153 to -1) were cloned into the pGL3 Basic vector (Addgene ##212936), resulting in the plasmid pGL3-APN-Luc. The pGL3-APN-Luc plasmid was methylated *in vitro* using CpG Methyltransferase M.SssI (NEB #M0226S). Briefly, 4 μg of plasmid was incubated with 4 μL CpG methyltransferase M.SssI, NEB buffer 2, and 640 μM S-adenosyl-methionine at 37°C. After 2 hours, an additional 640 μM S-adenosyl-methionine was added, and the reaction was incubated for another 2 hours at 37°C, followed by enzyme inactivation at 65°C for 20 min. Methylation status was verified by the EpiJET DNA Methylation Analysis Kit (MspI/HpaII) (Thermo #K1441). Unmethylated or methylated plasmids (1 μg) were digested with HpaII or MspI at 37°C for 1 hour and verified by 2% agarose gel electrophoresis.

For the luciferase reporter assay, HEK 293T cells grown in 24-well plates were transiently co-transfected with 500 ng of unmethylated or methylated luciferase reporter plasmid pGL3-APN-Luc and 50 of ng of pRL-TK plasmid (Promega # E2241). Luciferase activity was measured at 24 hours post-transfection using the Dual-Luciferase Reporter Assay System (Beyotime #RG027) and normalized to Renilla luciferase activity.

### Primary alveolar macrophage isolation

The procedure was performed as previously described^39^. Human alveolar macrophages (AMs) were isolated from bronchoalveolar lavage fluid (BALF). Samples were collected from young (n=4; 20, 22, 23, 25 years) and old (n=8; 53, 56, 61, 65, 67, 68, 76, 80 years) donors. BALF was obtained from routine bronchoscopies for noninflammatory/noninfectious disorders. The retrieved BALF was transferred immediately from the clinic to the laboratory at 4°C and processed by filtering through a cell strainer to remove mucus and particular debris. Alveolar macrophages were pelleted at 250 × g at 4°C for 10 min and seeded in 24-well plates for use. All procedures were conducted in accordance with the ethical standards of the Medical Ethics Council of Zhongshan Hospital (B2017-122). Informed consent was obtained from all participants.

### RNA extraction, reverse transcription, and qPCR

Total cellular RNA was extracted using TRIzol (Thermo #15596018) or RNAsimple Total RNA Kit (Tiangen #DP430) according to the manufacturer’s instructions. RNA was reverse transcribed into cDNA with the PrimeScript™ RT Reagent kit (Takara, RR047A) using a mix of Oligo dT primers and random 6-mers. qPCR was performed using the TB Green Premix Ex Taq™ II (Takara, RR820A) on a CFX Connect Real-Time System (Bio-Rad). Relative gene expression was calculated relative to GAPDH or human ACTB. All qPCR primer sequences are listed in **Supplementary Table 2**.

### RNA-seq analysis

Total cellular RNA of control and *UHRF1* sgRNA-edited A549-ACE2 cells was extracted using the RNAsimple Total RNA Kit (Tiangen #DP430) according to the manufacturer’s instructions. Library preparation was performed by Novogene, Inc., using at least 0.5 μg high-quality total RNA input per sample. mRNA enrichment was performed using poly-T oligo-attached magnetic beads. Sequencing libraries were generated using the NEBNext^®^ Ultra™ RNA Library Prep Kit for Illumina^®^ (NEB, USA) following the manufacturer’s recommendations. Libraries were sequenced on an Illumina Hiseq 2500 platform to generate paired-end reads.

Clean data (clean reads) were obtained by removing reads containing adapters, poly-N sequences, and low-quality reads from raw data. Differential expression analysis was performed using the DESeq2 R package (v1.36.0). P-values were adjusted using the Benjamini and Hochberg method to control the False Discovery Rate (FDR). Genes with an absolute Log_2_ fold change >2 and adjusted P-value <0.05 were considered as differentially expressed, and are listed in **Supplementary Table 3**. Volcano plots were generated in R using the EnhancedVolcano package. Gene Ontology (GO) enrichment and KEGG pathway analysis were performed using the ClusterProfiler 4.0 R package. Differential expression genes were extracted and used for functional clustering and network building using the STRING database and visualized in Cytoscape 3.10.1. The top 20 nodes ranked by MCC were calculated and graphed using the cytoHubba plugin.

### Focused CRISPR activation screen

A total of 2210 genes upregulated in *UHRF1*-knockout cells, identified by RNA-seq analysis, were selected for screening. A sub-libray of 8,676 activation sgRNAs targeting 2172 genes (approximately 4 sgRNAs per gene) was extracted from the human Calabrese CRISPR activation pooled library^63^ or custom-designed and synthesized (GENEWIZ). These sgRNAs were cloned into pXPR_502 (Addgene #96923). A549-ACE2-dCas9 cells were generated by transducing a packaged lentivirus derived from lenti dCAS-VP64_Blast (Addgene #61425). Cells were transduced with the sgRNA lentivirus sub-library and infected with HCoV-229E-mGreen or SARS-CoV-2 transcription- and replication-competent virus-like particles in which the N gene is replaced by the reporter GFP (trVLP-GFP)^38^ at an MOI of 0.5 for 24 h. Reporter-positive cells were sorted for genomic DNA extraction and sgRNA sequencing. The gene ranking was analyzed using MAGeCK software. sgRNA sequences and gene scores are listed in **Supplementary Table 4**.

### Protein 3D structure modeling

The 3D structures of PLAC1 (NP_001303816) and PLAC8 (NP_001124187) were predicted using AlphaFold2 via ColabFold v1.5.5 with default settings^69,70^. The models were visualized using PyMOL 2.3.2.

### Transciptomic data analysis

RNA-Seq data from 578 human lung samples, provided by the Genotype-Tissue Expression (GTEx) project, were reported as mean normalized transcripts per million (nTPM) and analyzed across three age groups: 20-39, 40-59, and 60-79. Correlation analyses were performed using the ggstatsplot R package. Single-cell RNA sequencing (scRNA-seq) datasets from lung samples of donors of varying ages were downloaded from a public database (GSE134355)^71^. Quality control was performed, and objects were generated from raw single-cell data using the Seurat package. Data normalization across sources was performed to mitigate systematic biases and ensure comparability. Two fetal (11-12 weeks), adult (21 years), and older (49 years) donors were analyzed. Cell types from all six donors were annotated using SingleR based on reference datasets, facilitating the identification and characterization of cell populations within the single-cell data.

### Statistical analysis

Statistical significance was assigned when P values were < 0.05 using Prism Version 9 (GraphPad). Data analysis was determined using ANOVA, unpaired t-test, or Mann-Whitney test, depending on data distribution and the number of comparison groups.

## DATA AVAILABILITY

The authors declare that all relevant data supporting the findings of this study are available within the paper and its Supplementary information. Raw RNA-sequencing data are deposited in SRA database and available (accession number, PRJNA1298838). Source Data are provided with this paper.

## ACKNOWLEDGEMENTS

Grants from the National Key Research and Development Program of China (2024YFC2607300 and 2020YFA0707701), National Natural Science Foundation of China (82341084, 32270163, 32041005), Program of Shanghai Academic Research Leader (22XD1420600), Shanghai Municipal Science and Technology Major Project (ZD2021CY001), Shenzhen Medical Research Fund (SMRF No. B2302029), Non-profit Central Research Institute Fund of Chinese Academy of Medical Sciences (2023-PT310-02) and China Postdoctoral Science Foundation (2023M740658) supported this work. We thank Jin Jin at the Center for Neuroimmunology and Health Longevity, the Third Affiliated Hospital of Sun Yat-sen University, for scientific and technical assistance. We wish to acknowledge Xiaoqing Sun, Yao Wang, and Shen Cai at Key Laboratory of Medical Molecular Virology (MOE/NHC/CAMS), Shanghai Frontiers Science Center of Pathogenic Microorganisms and Infection, School of Basic Medical Sciences of Fudan University for their help with next-generation sequencing, flow cytometry, and imaging analysis, respectively. We thank colleagues at the Biosafety Level 3 Laboratory of Fudan University for their technical assistance.

## AUTHOR CONTRIBUTIONS

P.W., Z.W., F.F., L.Y., Y.Z., C.L., and R.Z. performed the experiments. P.W., Z.W., F.F. and R.Z. designed the experiments. Y.Z., J.C., P.Z, S.Y, Y.H., J.Z., and C.L. provided technical or material support. P.Z., S.Y., Y.H., Q.D., Y.S., and R.Z. provided administrative and supervision support. P.W., Z.W., Z.G., and R.Z. performed data analysis. W.P. and R.Z. wrote the initial draft of the manuscript, with the other authors contributing to editing into the final form.

## COMPETING INTERESTS

The authors declare no competing interests.

## SUPPLEMENTARY FIGURES LEGENDS

**Supplementary Figure 1.**
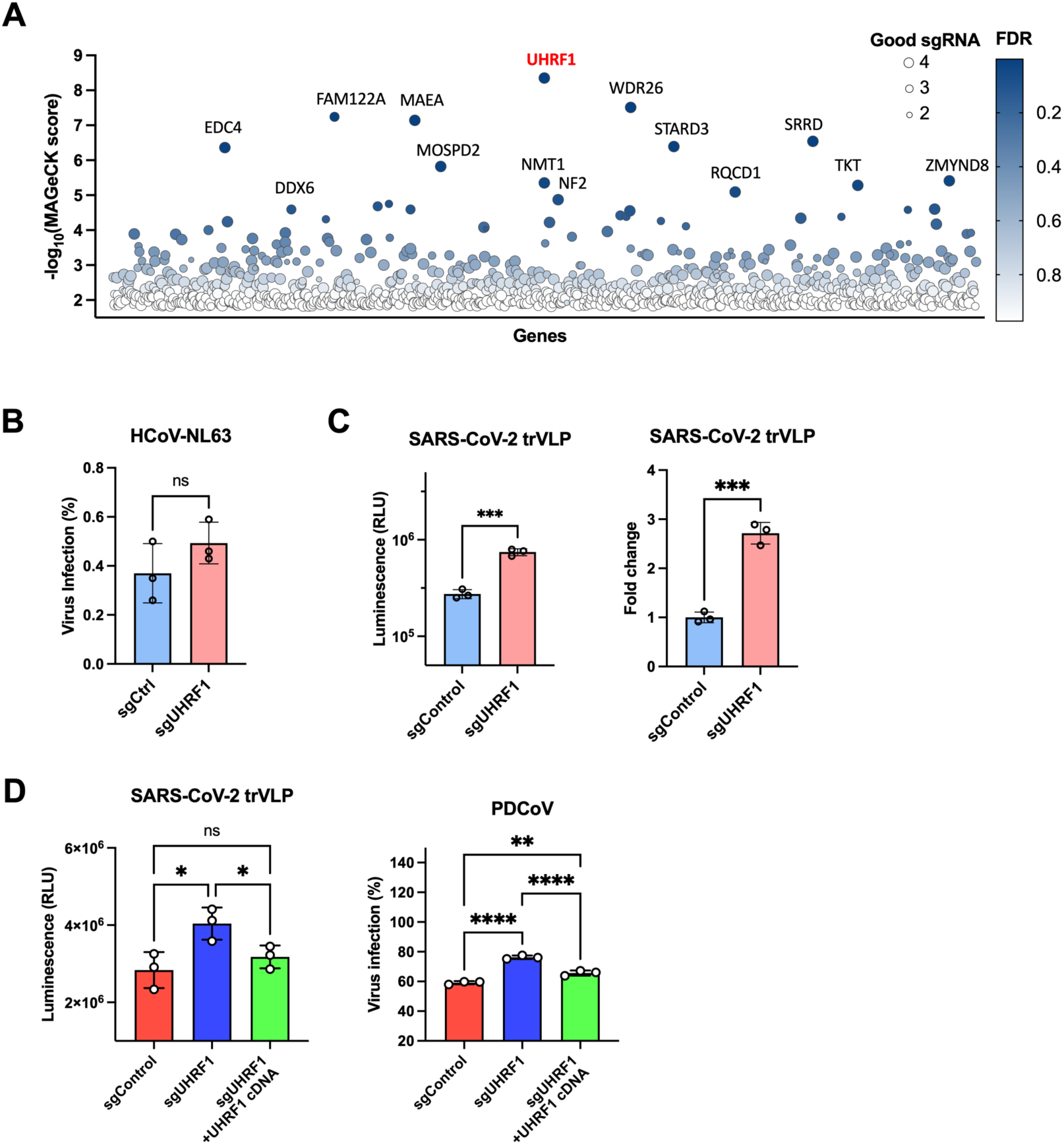
Genome-wide CRISPR knockout screen identifies top-ranked UHRF1 as a host restrict factor for HCoV-229E infection, and validation of UHRF1 for HCoV-NL63 and SARS-CoV-2. **A.** The A549 cell library containing genome-wide CRISPR knockout sgRNAs were infected with HCoV-229E-mGreen (MOI 0.5, 24 h). Infected reporter-positive cells were sorted for genomic DNA extraction and subsequent sgRNA sequence analysis. Genes were analyzed using MAGeCK software and ranked based on -log10 (MAGeCK score) and false discovery rate (FDR). The size of the circles represents the number of good sgRNAs enriched in the screen, while the color indicates the FDR. **B.** Control and *UHRF1*-knockout A549 cells without ectopic expression of ACE2 were infected with alphacoronavirus HCoV-NL63 (MOI 1, 24 h). Infection efficiency was determined by flow cytometry for the percentage of viral N protein-positive cells. **C.** Control and *UHRF1*-knockout A549 cells without ectopic expression of ACE2 were infected with betacoronavirus SARS-CoV-2 transcription- and replication-competent virus-like particles in which the N gene is replaced by reporter NanoLuc luciferase (trVLP) (MOI 0.5, 24 h). Infection efficiency was determined by measuring the luciferase activity, and the fold change was calculated by normalizing to the control. **D.** Infectivity of SARS-CoV-2 trVLP (MOI 0.5, 24 h) and PDCoV (MOI 0.3, 24 h) in *UHRF1*-knockout A549-ACE2 cells trans-complemented with UHRF1 cDNA. The luciferase activity or percentage of virus-positive cells were measured. Error bars represent standard deviations from three independent experiments. Unpaired t-test (B, C); one-way ANOVA with dunnett’s test (D). \**P* < 0.05; \*\**P* < 0.01; \*\*\**P* < 0.001; \*\*\*\**P* < 0.001; ns, not significant.

**Supplementary Figure 2.**
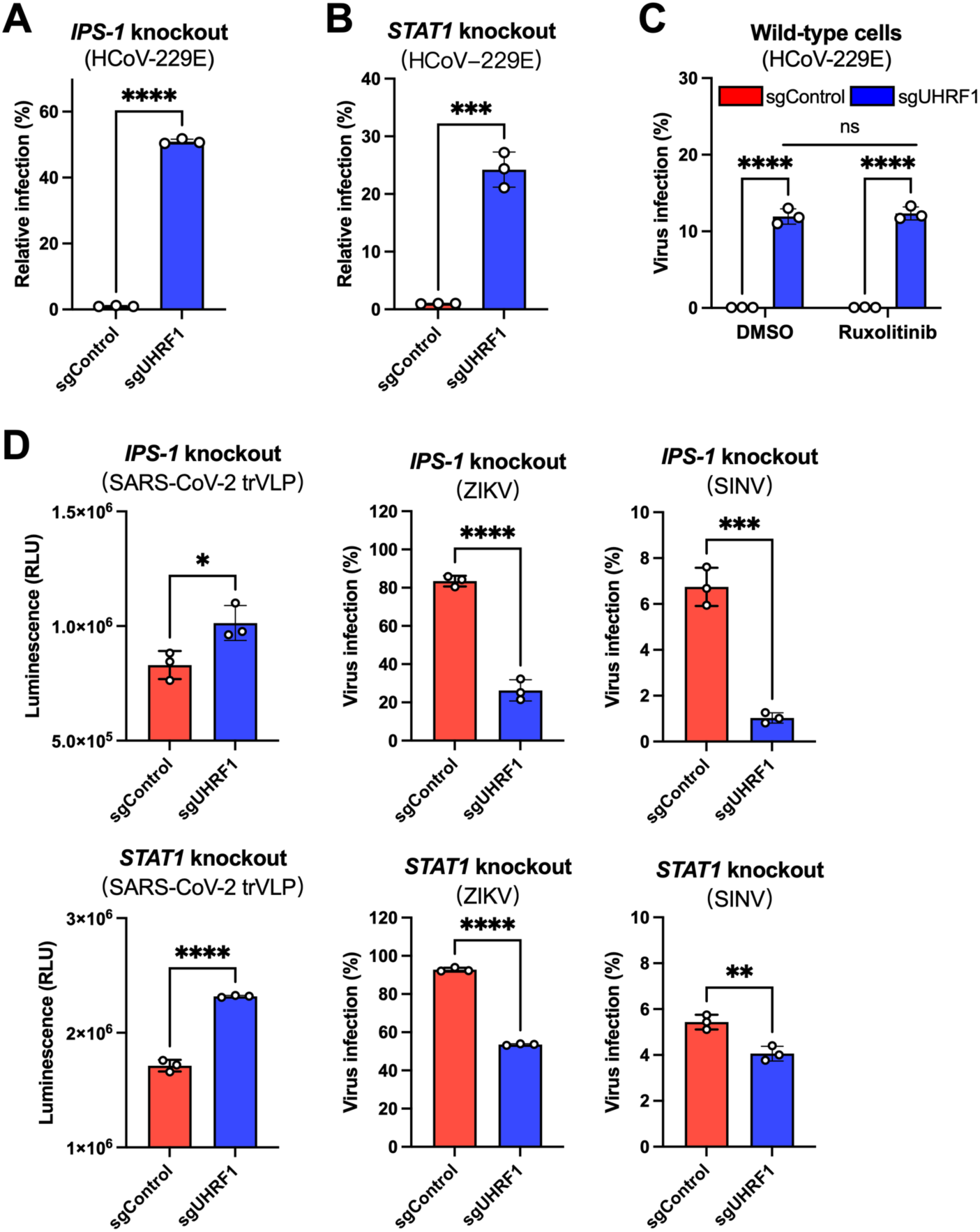
UHRF1 may restrict HCoV-229E and SARS-CoV-2 infection independently of IFN signaling pathway. **A-B.** Antiviral effect of UHRF1 on HCoV-229E infection in *IPS-1*- or *STAT1*-knockout A549 cells. Cells were infected with HCoV-229E (MOI 0.5, 24 h), and the percentage of N-positive cells was analyzed by flow cytometry. **C.** Control and *UHRF1*-knockout A549 cells were pre-treated with 5 μM Ruxolitinib for 1 h and then infected with HCoV-229E (MOI 0.5, 24 h) in the presence of the drug. The percentage of N-positive cells was analyzed by flow cytometry. **D.** Infectivity of SARS-CoV-2 trVLP (MOI 0.5, 24h), ZIKV (MOI 1, 24 h), and SINV (MOI 3, 24 h) in *IPS-1*- or *STAT1*-knockout A549-ACE2 cells edited with control or UHRF1 sgRNA. The luciferase activity or percentage of virus-positive cells were measured. Error bars represent standard deviations from three independent experiments. Unpaired t-test (A, B, D); two-way ANOVA with Sidak’s test (C). \**P* < 0.05; \*\**P* < 0.01; \*\*\**P* < 0.001; \*\*\*\**P* < 0.001; ns, not significant.

**Supplementary Figure 3.**
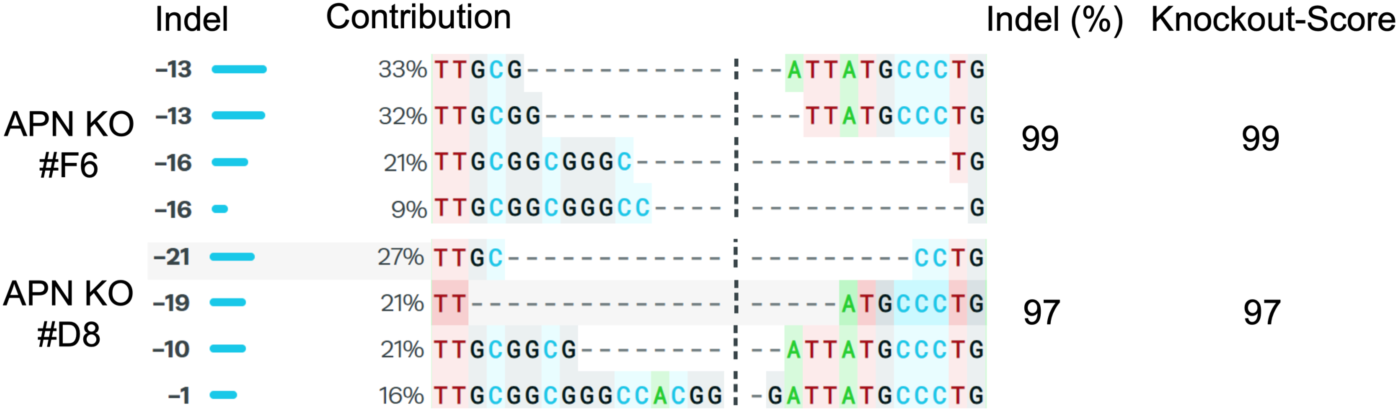
Inference of CRISPR Edits (ICE) analysis of *APN*-knockout clonal cell lines. A549 cells were edited by sgRNA targeting APN by lentivirus transduction, and clonal cell lines were obtained by limiting dilution. Single colonies were expanded for genomic DNA extraction. The sgRNA target regions were amplified, and knockout efficiency was determined by ICE analysis.

**Supplementary Figure 4.**
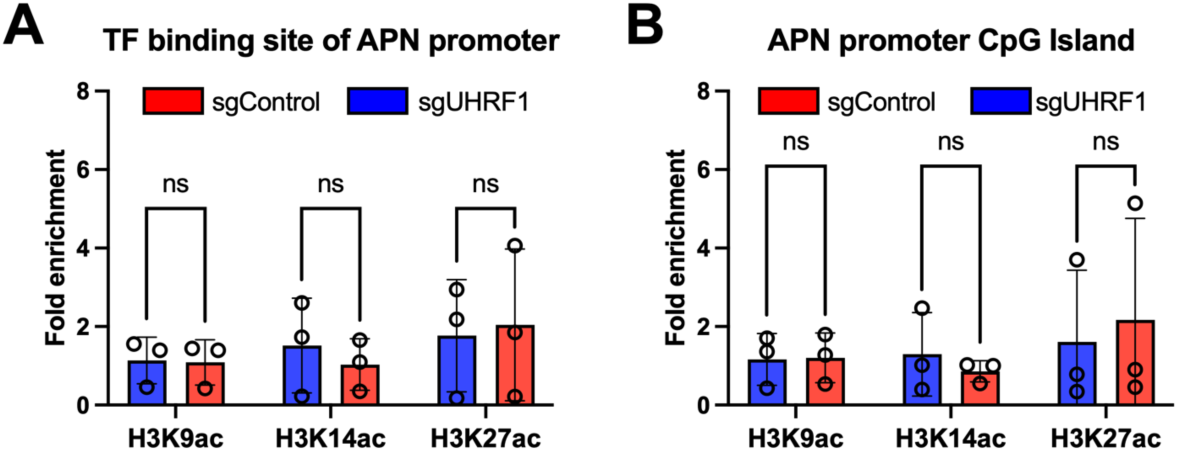
Knockout of UHRF1 does not affect the enrichment of acetylated histones on the APN promoter. **A-B.** Control and *UHRF1*-knockout A549 cells were collected for chromatin immunoprecipitation (ChIP)-qPCR. Antibodies against H3K9ac, H3K14ac, or H3K27ac were used to pull down the genomic DNA for qPCR targeting the transfection factor (TF) binding site or CpG island of APN proximal promoter. Error bars represent standard deviations from three independent experiments. Two-way ANOVA with Sidak’s test. ns, not significant.

**Supplementary Figure 5.**
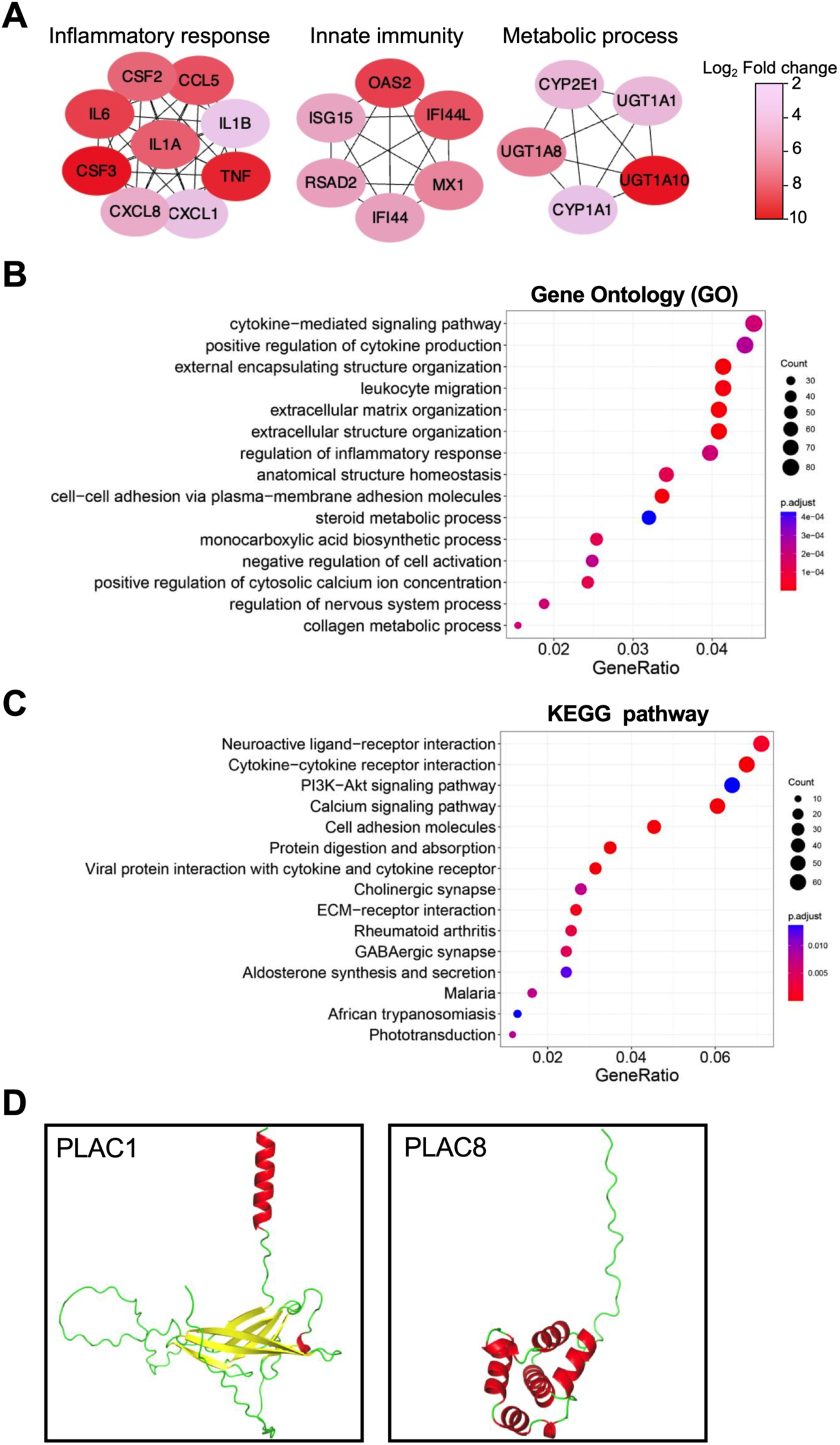
Differential expression of host genes between control and *UHRF1*-knockout cells, and structural comparison between PLAC1 and PLAC8. **A.** Network analysis of differentially expressed genes between control and *UHRF1*-knockout cells. Network analysis was conducted using the STRING database. The top 20 nodes ranked by MCC were calculated and graphed usng the cytoHubba plugin. The fold change of differentially expressed genes is indicated by color in the scale. **B-C.** Gene Ontology (GO) (biological process) (B) and KEGG pathway (C) enrichment analysis for differentially expressed genes. **D.** Structural comparison between PLAC1 and PLAC8 proteins. The 3D structures of PLAC1 (NP_001303816) (left) and PLAC8 (NP_001124187) (right) were predicted using AlphaFold2 via ColabFold v1.5.5 with default settings The models were visualized using PyMOL 2.3.2.

**Supplementary Figure 6.**
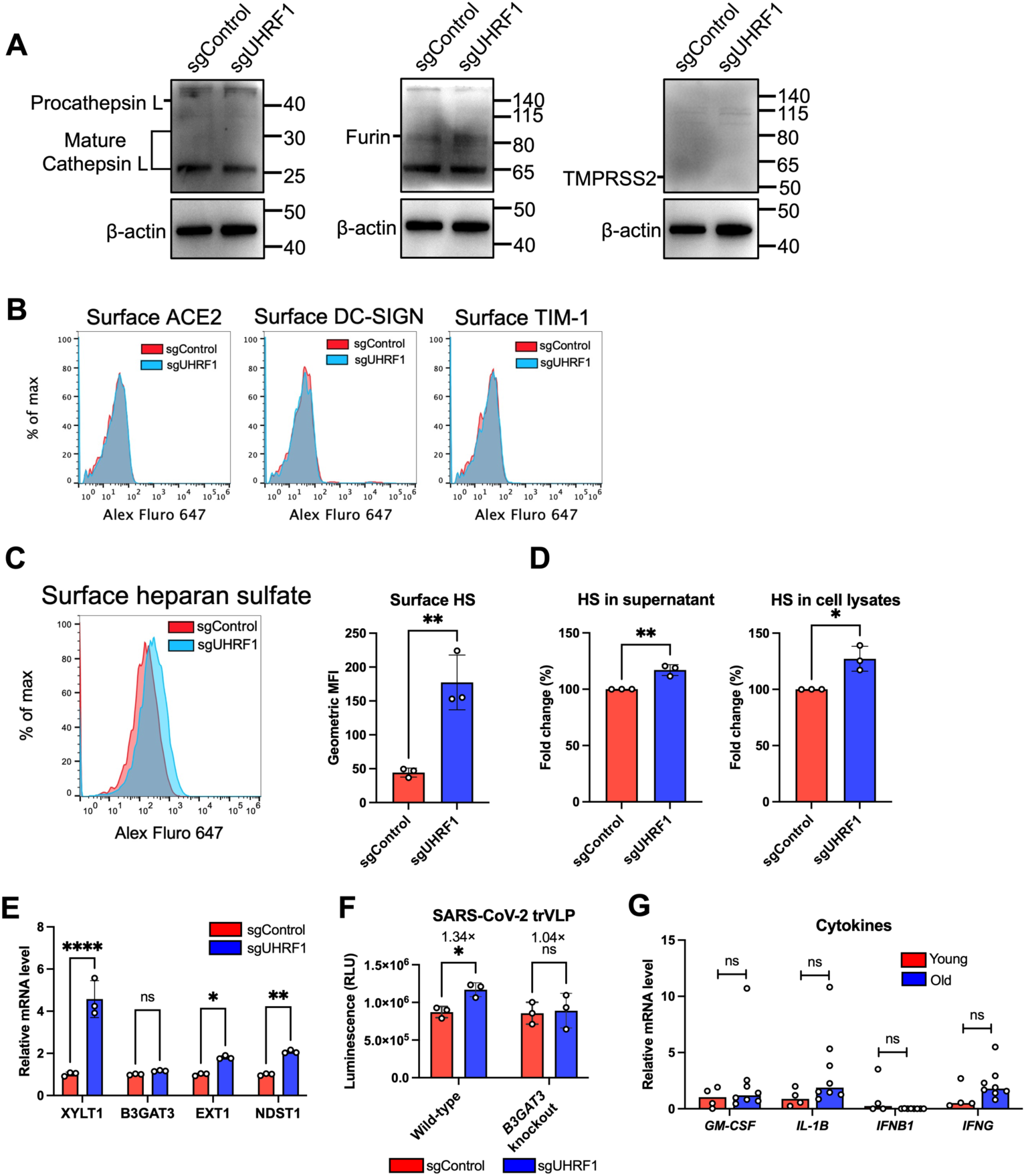
Effect of *UHRF1* knockout on the expression of coronavirus common entry factors, and expression of cytokines in HCoV-229E-infected alveolar macrophages. **A.** Western blotting analysis of CTSL, Furin, or TMPRSS2 in A549 cells edited with control or *UHRF1* sgRNA. **B.** Flow cytometry analysis of surface expression of ACE2, DC-SIGN, or TIM-1 in A549 cells edited with control or *UHRF1* sgRNA. **C.** Surface expression of heparan sulfate (HS) in A549 cells. Flow cytometry was performed to analyze the surface expression of heparan sulfate in A549 cells edited with control or *UHRF1* sgRNA (left). The geometric mean fluorescence intensity (MFI) of heparan sulfate labeling was quantified (right). **D.** Measurement of heparan sulfate expression by ELISA according to the manufacturer’s instructions (BBI, #D711289). ELISA was conducted to quantify heparan sulfate levels in the supernatants (left) and cell lysates (right) of A549 cells edited with control or *UHRF1* sgRNA. **E.** Expression of genes involved in heparan sulfate biosynthesis pathway. qRT-PCR was performed to detect the expression of key genes involved in heparan sulfate biosynthesis in A549 cells edited with control or *UHRF1* sgRNA. **F.** Infection of SARS-CoV-2 trVLP (MOI 0.5, 24 h) in wild-type or *B3GAT3*-knckout A549 cells edited with control or *UHRF1* sgRNA. The NanoLuc luciferase activity expressed by the virus was measured. **G.** Expression of cytokine genes in alveolar macrophages from young (n=4) and old (n=8) donors infected with HCoV-229E. Virus-infected cells were harvested at 24 h post-infection, and total cellular RNA was extracted for qRT-PCR targeting the indicated genes. Error bars represent standard deviations (C-F); the median values are shown (G). Unpaired t test (C-E); two-way ANOVA with Sidak’s test (F). Mann-Whitney test (G). *P < 0.05; **P < 0.01; ****P < 0.001; ns, not significant.

## SUPPLEMENTARY TABLE LEGENDS

Supplementary Table 1. List of genes and scores after MAGeCK analysis of genome-wide knockout screens for HCoV-229E (see Excel file).

Supplementary Table 2. sgRNA sequences of genes selected for validation and other editing experiments, and primer sequences for PCR or qRT-PCR experiments (see Excel file).

Supplementary Table 3. List of differentially expressed genes from A549 cells edited with control or *UHRF1* sgRNA (see Excel file).

Supplementary Table 4. List of genes and the relevant sgRNA sequences selected for focused sub-library construction, and scores after MAGeCK analysis of focused activation screens for HCoV-229E and SARS-CoV-2 (see Excel file).

## Notes

### Competing Interest Statement

The authors have declared no competing interest.

## REFERENCES

1. Lamers, M.M. & Haagmans, B.L. SARS-CoV-2 pathogenesis. Nat Rev Microbiol 20, 270–284 (2022).

2. V’Kovski, P., Kratzel, A., Steiner, S., Stalder, H. & Thiel, V. Coronavirus biology and replication: implications for SARS-CoV-2. Nat Rev Microbiol 19, 155–170 (2021).

3. Tyrrell, D.A., Cohen, S. & Schlarb, J.E. Signs and symptoms in common colds. Epidemiol Infect 111, 143–56 (1993).

4. Pene, F. et al. Coronavirus 229E-related pneumonia in immunocompromised patients. Clin Infect Dis 37, 929–32 (2003).

5. Sun, W. et al. A severe case of human coronavirus 229E pneumonia in an elderly man with diabetes mellitus: a case report. BMC Infect Dis 21, 524 (2021).

6. Villamil-Gomez, W.E. et al. Fatal human coronavirus 229E (HCoV-229E) and RSV-Related pneumonia in an AIDS patient from Colombia. Travel Med Infect Dis 36, 101573 (2020).

7. Funk, C.J. et al. Infection of human alveolar macrophages by human coronavirus strain 229E. J Gen Virol 93, 494–503 (2012).

8. Desforges, M., Miletti, T.C., Gagnon, M. & Talbot, P.J. Activation of human monocytes after infection by human coronavirus 229E. Virus Res 130, 228–40 (2007).

9. Dominguez, S.R., Travanty, E.A., Qian, Z. & Mason, R.J. Human coronavirus HKU1 infection of primary human type II alveolar epithelial cells: cytopathic effects and innate immune response. PLoS One 8, e70129 (2013).

10. Wang, Z., Li, S. & Huang, B. Alveolar macrophages: Achilles’ heel of SARS-CoV-2 infection. Signal Transduct Target Ther 7, 242 (2022).

11. Malainou, C., Abdin, S.M., Lachmann, N., Matt, U. & Herold, S. Alveolar macrophages in tissue homeostasis, inflammation, and infection: evolving concepts of therapeutic targeting. J Clin Invest 133(2023).

12. Yeager, C.L. et al. Human aminopeptidase N is a receptor for human coronavirus 229E. Nature 357, 420–2 (1992).

13. Bostick, M. et al. UHRF1 plays a role in maintaining DNA methylation in mammalian cells. Science 317, 1760–4 (2007).

14. Wu, Y. et al. UHRF1 establishes crosstalk between somatic and germ cells in male reproduction. Cell Death Dis 13, 377 (2022).

15. Kostyrko, K. et al. UHRF1 is a mediator of KRAS driven oncogenesis in lung adenocarcinoma. Nat Commun 14, 3966 (2023).

16. Hu, C.L. et al. Targeting UHRF1-SAP30-MXD4 axis for leukemia initiating cell eradication in myeloid leukemia. Cell Res 32, 1105–1123 (2022).

17. Xu, X. et al. Nuclear UHRF1 is a gate-keeper of cellular AMPK activity and function. Cell Res 32, 54–71 (2022).

18. Gao, Z.J. et al. Single-nucleotide methylation specifically represses type I interferon in antiviral innate immunity. J Exp Med 218(2021).

19. Wang, M. et al. UHRF1 Deficiency Inhibits Alphaherpesvirus through Inducing RIG-I-IRF3-Mediated Interferon Production. J Virol 97, e0013423 (2023).

20. Liang, T. et al. UHRF1 Suppresses HIV-1 Transcription and Promotes HIV-1 Latency by Competing with p-TEFb for Ubiquitination-Proteasomal Degradation of Tat. mBio 12, e0162521 (2021).

21. Verdikt, R. et al. Novel role of UHRF1 in the epigenetic repression of the latent HIV-1. EBioMedicine 79, 103985 (2022).

22. Guo, R. et al. DNA methylation enzymes and PRC1 restrict B-cell Epstein-Barr virus oncoprotein expression. Nat Microbiol 5, 1051–1063 (2020).

23. Schneider, W.M. et al. Genome-Scale Identification of SARS-CoV-2 and Pan-coronavirus Host Factor Networks. Cell 184, 120–132 e14 (2021).

24. Irwin, R.E. et al. The UHRF1 protein is a key regulator of retrotransposable elements and innate immune response to viral RNA in human cells. Epigenetics 18, 2216005 (2023).

25. Conant, D. et al. Inference of CRISPR Edits from Sanger Trace Data. CRISPR J 5, 123–130 (2022).

26. Guan, D., Factor, D., Liu, Y., Wang, Z. & Kao, H.Y. The epigenetic regulator UHRF1 promotes ubiquitination-mediated degradation of the tumor-suppressor protein promyelocytic leukemia protein. Oncogene 32, 3819–28 (2013).

27. Oh, Y.M. et al. Epigenetic regulator UHRF1 inactivates REST and growth suppressor gene expression via DNA methylation to promote axon regeneration. Proc Natl Acad Sci U S A 115, E12417–E12426 (2018).

28. Kim, M.J. et al. UHRF1 Induces Methylation of the TXNIP Promoter and Down-Regulates Gene Expression in Cervical Cancer. Mol Cells 44, 146–159 (2021).

29. Christman, J.K. 5-Azacytidine and 5-aza-2’-deoxycytidine as inhibitors of DNA methylation: mechanistic studies and their implications for cancer therapy. Oncogene 21, 5483–95 (2002).

30. Kratzel, A. et al. A genome-wide CRISPR screen identifies interactors of the autophagy pathway as conserved coronavirus targets. PLoS Biol 19, e3001490 (2021).

31. Zhang, Z.M. et al. Structural basis for DNMT3A-mediated de novo DNA methylation. Nature 554, 387–391 (2018).

32. Mahoney, K.M., Petrovic, N., Schacke, W. & Shapiro, L.H. CD13/APN transcription is regulated by the proto-oncogene c-Maf via an atypical response element. Gene 403, 178–87 (2007).

33. Kong, X. et al. Defining UHRF1 Domains that Support Maintenance of Human Colon Cancer DNA Methylation and Oncogenic Properties. Cancer Cell 35, 633–648 e7 (2019).

34. Li, T. et al. Structural and mechanistic insights into UHRF1-mediated DNMT1 activation in the maintenance DNA methylation. Nucleic Acids Res 46, 3218–3231 (2018).

35. Zhao, Q. et al. Dissecting the precise role of H3K9 methylation in crosstalk with DNA maintenance methylation in mammals. Nat Commun 7, 12464 (2016).

36. Szklarczyk, D. et al. The STRING database in 2023: protein-protein association networks and functional enrichment analyses for any sequenced genome of interest. Nucleic Acids Res 51, D638–D646 (2023).

37. Chin, C.H. et al. cytoHubba: identifying hub objects and sub-networks from complex interactome. BMC Syst Biol 8 Suppl 4, S11 (2014).

38. Ju, X. et al. A novel cell culture system modeling the SARS-CoV-2 life cycle. PLoS Pathog 17, e1009439 (2021).

39. Feng, F. et al. A CRISPR activation screen identifies genes that enhance SARS-CoV-2 infection. Protein Cell 14, 64–68 (2023).

40. Hudak, A., Letoha, A., Szilak, L. & Letoha, T. Contribution of Syndecans to the Cellular Entry of SARS-CoV-2. Int J Mol Sci 22(2021).

41. Ugalde, A.P. et al. Autophagy-linked plasma and lysosomal membrane protein PLAC8 is a key host factor for SARS-CoV-2 entry into human cells. EMBO J 41, e110727 (2022).

42. Tse, L.V. et al. Genomewide CRISPR knockout screen identified PLAC8 as an essential factor for SADS-CoVs infection. Proc Natl Acad Sci U S A 119, e2118126119 (2022).

43. Millet, J.K., Jaimes, J.A. & Whittaker, G.R. Molecular diversity of coronavirus host cell entry receptors. FEMS Microbiol Rev 45(2021).

44. Zhu, X. et al. Porcine Deltacoronavirus nsp5 Antagonizes Type I Interferon Signaling by Cleaving STAT2. J Virol 91(2017).

45. Zheng, Y. et al. SARS-CoV-2 NSP5 and N protein counteract the RIG-I signaling pathway by suppressing the formation of stress granules. Signal Transduct Target Ther 7, 22 (2022).

46. Liu, G. et al. ISG15-dependent activation of the sensor MDA5 is antagonized by the SARS-CoV-2 papain-like protease to evade host innate immunity. Nat Microbiol 6, 467–478 (2021).

47. Li, W. et al. Broad receptor engagement of an emerging global coronavirus may potentiate its diverse cross-species transmissibility. Proc Natl Acad Sci U S A 115, E5135–E5143 (2018).

48. Milewska, A. et al. Human coronavirus NL63 utilizes heparan sulfate proteoglycans for attachment to target cells. J Virol 88, 13221–30 (2014).

49. Madu, I.G. et al. Heparan sulfate is a selective attachment factor for the avian coronavirus infectious bronchitis virus Beaudette. Avian Dis 51, 45–51 (2007).

50. LeBlanc, E.V. & Colpitts, C.C. The green tea catechin EGCG provides proof-of-concept for a pan-coronavirus attachment inhibitor. Sci Rep 12, 12899 (2022).

51. Fuochi, V. et al. Heparan Sulfate and Enoxaparin Interact at the Interface of the Spike Protein of HCoV-229E but Not with HCoV-OC43. Viruses 15(2023).

52. Clausen, T.M. et al. SARS-CoV-2 Infection Depends on Cellular Heparan Sulfate and ACE2. Cell 183, 1043–1057 e15 (2020).

53. Jones, P.A. Functions of DNA methylation: islands, start sites, gene bodies and beyond. Nat Rev Genet 13, 484–92 (2012).

54. Cedar, H. & Bergman, Y. Linking DNA methylation and histone modification: patterns and paradigms. Nat Rev Genet 10, 295–304 (2009).

55. Bird, A. DNA methylation patterns and epigenetic memory. Genes Dev 16, 6–21 (2002).

56. Rebendenne, A. et al. Bidirectional genome-wide CRISPR screens reveal host factors regulating SARS-CoV-2, MERS-CoV and seasonal HCoVs. Nat Genet 54, 1090–1102 (2022).

57. Biering, S.B. et al. Genome-wide bidirectional CRISPR screens identify mucins as host factors modulating SARS-CoV-2 infection. Nat Genet 54, 1078–1089 (2022).

58. Baggen, J. et al. Genome-wide CRISPR screening identifies TMEM106B as a proviral host factor for SARS-CoV-2. Nat Genet (2021).

59. Wei, J. et al. Genome-wide CRISPR Screens Reveal Host Factors Critical for SARS-CoV-2 Infection. Cell (2020).

60. Wang, R. et al. Genetic Screens Identify Host Factors for SARS-CoV-2 and Common Cold Coronaviruses. Cell (2020).

61. Daniloski, Z. et al. Identification of Required Host Factors for SARS-CoV-2 Infection in Human Cells. Cell (2020).

62. Zhu, Y. et al. A genome-wide CRISPR screen identifies host factors that regulate SARS-CoV-2 entry. Nat Commun 12, 961 (2021).

63. Doench, J.G. et al. Optimized sgRNA design to maximize activity and minimize off-target effects of CRISPR-Cas9. Nat Biotechnol 34, 184–191 (2016).

64. Feng, F. et al. DAZAP2 functions as a pan-coronavirus restriction factor by inhibiting viral entry and genomic replication. *bioRxiv*, 2025.02.04.636569 (2025).

65. Yang, Q. et al. Farnesyltransferase inhibitor lonafarnib suppresses respiratory syncytial virus infection by blocking conformational change of fusion glycoprotein. Signal Transduct Target Ther 9, 144 (2024).

66. Sun, W. et al. Cross-species infection potential of avian influenza H13 viruses isolated from wild aquatic birds to poultry and mammals. Emerg Microbes Infect 12, e2184177 (2023).

67. Ma, X. et al. The single amino acid change of R516K enables efficient generation of vesicular stomatitis virus-based Crimean-Congo hemorrhagic fever reporter virus. *bioRxiv*, 2025.01.05.631422 (2025).

68. Wang, J. et al. The mycobacterial phosphatase PtpA regulates the expression of host genes and promotes cell proliferation. Nat Commun 8, 244 (2017).

69. Mirdita, M. et al. ColabFold: making protein folding accessible to all. Nat Methods 19, 679–682 (2022).

70. Jumper, J. et al. Highly accurate protein structure prediction with AlphaFold. Nature 596, 583–589 (2021).

71. Han, X. et al. Construction of a human cell landscape at single-cell level. Nature 581, 303–309 (2020).

